# Development of low cost, robust and reproducible biofilm static and pharmacodynamic assays

**DOI:** 10.64898/2026.07.27.740918

**Authors:** Marie Attwood, Pippa Griffin, Alasdair MacGowan, Shona Nelson, Alan Noel, Patryk Smorowinski, Dann Turner

## Abstract

**Background:** The complexity of diagnosing and treating biofilm-associated infections necessitates a comprehensive strategy to mitigate the rising rates of antimicrobial resistance (AMR). Microtiter plate methods are used globally for determination of biofilm eradication concentrations (MBEC) but few have been adapted to observe pharmacodynamic observations. Here, we describe a method which allows for both static and pharmacodynamic assays of biofilm evaluation.

**Methods:** A total of 150 clinical isolates from Southmead Hospital were assessed, representing five bacterial species (N=30 per bacterial species): *Pseudomonas aeruginosa, Escherichia coli, Streptococcus pneumoniae, Staphylococcus aureus* and *Klebsiella pneumoniae*. MBECs were determined using a developed method using 96 well plates and glass beads. MBECs of seven different antibiotics were compared to those determined using the established Calgary biofilm device (CBD). Dynamic pharmacodynamic evaluations to produce Biofilm Time Kill curve (BTKC) based on published planktonic time kill curve (TKC) data and ISO recommendations were carried out using the glass bead model for *K. pneumoniae* and ciprofloxacin, *S. aureus* and levofloxacin and *S. pneumoniae* and vancomycin. Quantification of biofilm biomass was assessed at 0, 2, 4, 8 and 24 hours and compared to planktonic culture survival under comparable challenge conditions.

**Results:** Comparing MBEC results for all bacterial strains and antibiotic challenges showed no statistical difference between the glass bead and CBD methods (P <0.05). Biofilm BTKC AUBKC were inferior to planktonic equivalents but demonstrated specific pharmacodynamic patterns of biofilm reduction efficacy. MBEC correlated with biofilm BTKC penetration in line with clinical observations for *S. aureus* vs vancomycin and *S. pneumoniae* vs levofloxacin.

**Conclusions:** The glass bead biofilm models provide robust, reproducible alternatives to the traditional methods of determining MBEC and bridge the gap with biofilm pharmacodynamic evaluations. These methods also provide a low-cost option to current methods as only standard laboratory equipment is required, allowing for the generation of comprehensive data sets. This ensures greater translatability to complex *in vitro* models and clinical scenarios.

## Introduction

Biofilms are surface-associated, complex, microbial communities ^1^ commonly comprised of a range of prokaryotic and/or eukaryotic cells, embedded within a matrix of extracellular polymeric substances (EPS). As biofilms mature, their heterogeneity increases and this varied, dynamic lifestyle poses a challenge to effective antimicrobial chemotherapy. Microbial biofilm communities play a significant role in the most chronic infection often resulting in therapeutic failure^2^. With the increasing urgency for global initiatives to prevent escalating Antimicrobial Resistance (AMR) levels, there is an urgent need for the development of novel anti-bacterial or combination antibiotic therapies. Previous novel or combination therapies^3^,6,^4^ ^5^ have focused on the eradication or reduction of bacterial burden with the aim to prevent regrowth of a bacterial population and therefore reduce the risk of emergence of resistance. However, biofilm-forming abilities of bacterial populations are not routinely studied in diagnostic laboratories, a limitation that could be attributed to a lack of cost-appropriate and/or reproducible international methodologies.

To date there are an array of techniques available to evaluate biofilms, each offering distinct advantages such as high throughput or low cost, but also presenting limitations, including the requirement for specialised material. They can be divided into four defined categories; direct measurements, indirect measurement, *in situ* and *ex situ* methodologies^6^.

This current study is based on *ex situ* techniques which can determine biofilm minimum eradication points that can be adapted/used to enable pharmacodynamic assessments over time.

Microtiter plates are perhaps the most common method used globally for the determination of biofilm eradication concentrations by growing bacterial cells statically on polystyrene microtiter plate wells^7^. Whilst this high throughput and low-cost method is useful for screening anti-biofilm compounds, limitations remain. This method is based on non-specific staining with no differentiation between live and dead cells, is impacted by stained sedimented cells and endpoint-only results hinder studies of chronic biofilm or planktonic-biofilm interactions. These drawbacks restrict its use to short term simulations, with known variable reproducibility and a lack of translatability to more complex *in vitro* systems^8^.

The Calgary Biofilm Device (CBD) addresses reproducibility issues caused by cell sedimentation when using staining techniques. CBD device consists of a two-part reaction vessel with 96 pegs that create consistent shear force for biofilm formation^9^. The resulting biofilm can be evaluated by CFU enumeration after peg removal and sonication^10^ which offers a high-throughput screening method with much improved reproducibility. However, like microtiter plates, CBD also has limitations: endpoint only measurements (restricting studies to early-stage biofilm interactions), short term simulations due to nutrient depletion (<72 hours), toxic by-product accumulation and evaporation of media within the 96 well plate. Variability from technical factors i.e. inconsistent peg removal or differences in sonication placement can alter recovery rates and, in turn, affect reproducibility^11^. Peg lids within CBD are typically made of specific materials, however, there are a range of coatings available which aims to address any non-specific binding or hydrophobic compound related issues. Despite the vast improvement in this biofilm screening device, the primary drawback of this device remains its relative expense^12^.

The FlexiPeg (FP) is a 3D-printed alternative to the CBD, designed to provide enhanced versatility. Specifically; easier single peg removal, 3-D pegs design, autoclavable coatings and customizable material by printing allowing the accurate mimicking of a medical surface device. FP represents a highly diverse biofilm screening and potential useful biofilm TKC (BTKC) device, although this model would require adjustments for broader applications^12,13^.

There are other biofilm related products and methodologies, for example the biofilm ring test. The principal of this method is that the presence of a biofilm prevents the movement of paramagnetic beads when exposed to a magnetic field^14^. Since no medium intervention is needed after inoculation, it reduces inter laboratory variation and enables repeated measurement, generating biofilm formation data and provides information which may compliment classical antibiograms. However, this technique requires specialised equipment, reagents, and reproducibility is impacted by sedimented cells, which accumulate proportionately as the biofilm develops^15^.

The ideal techniques for the evaluation of novel antibiofilm compounds would include companion *in vitro* biofilm MBEC and BTKC pharmacodynamic simulations where the methodologies are readily incorporated into a dynamic *in vitro* modelling system. All these methodologies must have equal consideration for versatility, high throughput, reproducibility, and affordability. Key aspects include consistency in the production of replicate biofilms, removal of planktonic debris, effective biofilm cell recovery and enumeration and multiple biofilm measurements over the course of experimental simulations. These specific factors could then be applied in more traditional *in vitro* modelling systems (where human pharmacokinetic antibiotic levels can be mimicked) such as dilutional models which simulate human bloodstream infections or neutropenic thigh models to assist in the establishment of optimised dosing regimens^16–18^. This approach could enable the development of site-specific models that replicate specific but dynamic environments; bladder *in vitro* models simulating voiding, reduction of total culture volume for pleural effusion studies, limited nutrient availability for bone joint infections.

We propose the development of a low cost, user friendly, robust and reproducible high throughput screening biofilm assay (referred to as NBT). This assay uses shear force based, surface-attached biofilm methodology^19^ comprising of 96 well plates and glass beads. This is designed to enable initial MBEC evaluations, which support preliminary dynamic testing (BTKC) and allows integration into more complex pharmacokinetic and pharmacodynamic *in vitro* experiments.

## Materials

### Bacterial strains

Biofilm eradication concentration: For the determination of minimum biofilm eradication concentration a total of 150 clinical isolates collected from patients at Southmead Hospital, North Bristol NHS Trust were assessed. Clinical isolates comprised 30 independent cultures each of *Pseudomonas aeruginosa, Escherichia coli, Streptococcus pneumoniae, Staphylococcus aureus* and *Klebsiella pneumoniae.* Type strains of each species were also assessed, including *P. aeruginosa* ATCC 27853 (PSEAE), *E. coli* ATCC 25922 (ESCCO), *S. pneumoniae* ATCC 49619 (STRPN), *S. aureus* ATCC 29213 (STAAU) and *K. pneumoniae* NCTC 700603 (KLEPN).

BTKC evaluations were performed to cover gram positive, negative and microaerophilic organisms and used KLEPN 700603, STAAU 29213 and STRPN 49619.

### Media

Mueller Hinton broth II (BD, product code 212322) was used for all experiments. Nutrient agar plates (Oxoid, UK, product reference PO0155A) were used to recover bacterial strains from storage and for inoculum confirmations.

### Antibiotics

Bacterial isolates were tested against with the following antibiotics (Table 1.); Vancomycin, Ciprofloxacin, Colistin, Linezolid, Levofloxacin, Piperacillin and Tazobactam (Merck Life Science UK limited). Stock and working solutions of antibiotics were prepared in line with CLSI and EUCAST (Version 5.0 Jan 2024) guidelines.

**Table 1.** Bacterial strain and antibiotic biofilm combinations.

| Bacterial strain | Antibiotics biofilm combinations |
| --- | --- |
| STAAU | Vancomycin<br>Ciprofloxacin |
| PSEAE | Ciprofloxacin<br>Colistin |
| STRPN | Linezolid<br>Levofloxacin |
| ESCCO | Ciprofloxacin<br>Piperacillin/tazobactam |
| KLEPN | Ciprofloxacin<br>Colistin |

## Methods

### MIC determination

MIC determinations were performed by broth-microdilution in strict accordance with ISO 7218:2024 and with European Committee on Antimicrobial Susceptibility Testing (EUCAST) guidelines that aim to assess the susceptibility of non-fastidious organisms to antimicrobial agents. (V 5.0, 1^st^ of January 2024). Anaerobic conditions were used for *S. pneumoniae* in accordance with CLSI guidelines (M11-M56). Planktonic MIC ranges and MBEC were determined using standard doubling dilution format. Quality control strains (ATCC strains) were included per run with a total of six replicates.

### Growth subcultures and inoculum preparations – CBD and NBT

All bacterial strains used were sub-cultured from glycerol stocks stored at -80°C onto non-selective nutrient agar. Cultures were incubated at 37°C for 18-24 hours prior to associated experimental procedures. Inoculations were prepared in MHB to achieve turbidity equivalent to a 0.5 McFarland standard by eye or, alternatively, using a spectrophotometer at a wavelength of 625nm, (range 0.08-0.13) in accordance with ISO standards. A target cell density for CBD and NBT MBEC of 5.5x10^5^

CFU/mL (as per manufacturer’s instructions), was employed. Inoculum checks were performed by serial diluting and spot plating the diluted culture to confirm cell density per well.

### CBD (Calgary Biofilm Device MBEC®) Assay - Set up

A 200 µL volume of inoculum was transferred into each testing well of a CBD kit (Innovotech INC, Calgary, Canada) before placing the peg lid onto the base. The assembly was agitated according to manufacturer’s instruction, using a rocking table, and incubated for a minimum of 16 hours. Longer incubation periods were used to assess more developed biofilm formation, with acute biofilm formation up to and including 48 hours. Incubation of greater than 48 hours represents more mature/chronic biofilms but can be affected by evaporation of media.

### CBD (Calgary Device MBEC®) Assay – challenge plate

A standard 96 well dilution series with relevant antibiotic, including Growth control (GC) and Sterility control (SC) wells were organised which can be seen in Supplementary Figure 1

### CBD (Calgary Device MBEC®) Assay - Biofilm growth check

After 48 hours of growth, planktonic bacteria were removed by transferring the peg lid into 96-well microtitre plates containing 200 µL of sterile saline, ensuring that pegs were immersed for 10 seconds (wash plate). After rinsing, pegs were removed from the lid, placed into 200 µL recovery media (Mueller Hinton broth II) and sonicated for 30 ±5 minutes (medium Aquasonic model 250HT, VWR scientific) to dislodge and disperse the biofilm. Enumeration was performed using a spiral plater (Don Whitley Scientific, UK) onto non-selective agar. Following incubation (18-20 hours at 37°c) biofilm densities were expressed as CFU/mL.

### CBD (Calgary Device MBEC®) Assay - Determination of Minimum biofilm eradication concentration

After biofilm growth check determination (48 hours), CBD MBECs were performed as stated. Peg lids were immediately transferred to the antibiotic challenge plates after rinsing and incubated for a further 24 hours at 37 °C. After incubation, biofilms were disrupted by sonication and then checked visually for turbidity. The MBEC was assigned to the lowest antibiotic concentration where no visible turbidity was observed.

### NBT Biofilm determination assay

For the NBT biofilm assay, 200 µL of inoculum was transferred to the wells of a standard (v-bottom) microtiter plate (Alpha labs) each containing a single sterile 3 mm glass bead. The microtiter plate was agitated continuously using a rocking platform and incubated for 48 hours for biofilm generation. Rinsing, turbidity evaluation and determination of the MBEC was performed in the same way as described for the CBD.

**Figure 2.**
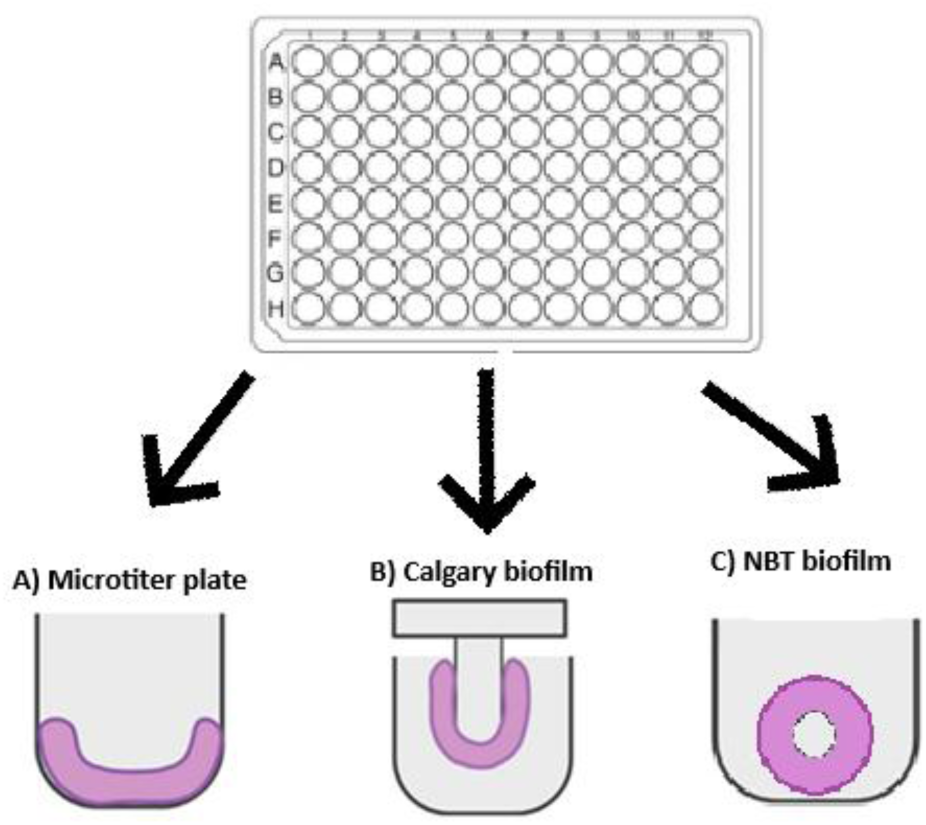
CBD and NBT Biofilm MBEC determination.

### Establishing initial biofilm for pharmacodynamic interaction in 10mL culture vessels

To establish larger biofilms suitable for pharmacodynamic interactions, 20 glass beads were added to 10 mL of MHB in glass universals and incubated for a minimum of 1 hour prior to bacterial inoculation, ensuring all material had reached 37.5°C. Relevant bacteria were inoculated into the culture vessels containing glass beads (as previously described above). Target starting bacterial suspensions for BTKC was equivalent to 1.5x10^6^ CFU/mL, which is consistent with in vivo TKC studies targets^20^. To obtain a biofilm density of approximately 1.5x10^6^ CFU/mL, the culture vessels were incubated at 37°C, with shaking at 150 RPM, for 48 hours.

### Biofilm Time Kill Curve

*The protocol for biofilm TKC is based on well-known published data which has been used successfully at NBT for the progression of novel molecules through drug development platforms*^21^*. (Ref). Standard planktonic TKCs were performed for each bacterial strain for comparison. Slight alterations biofilm TKC simulations are stated below*.

Once biofilm had developed as described previously and biofilm baseline CFU/mL was confirmed, all available media was removed from the universal to ensure biofilm-only evaluations and was replaced by 10mL fresh MHBII. If simultaneous planktonic and biofilm CFU/mL evaluations were required, then MHBII media was not replaced, and planktonic CFU/mL was also determined. Antibiotic concentrations established in MHBII in MIC multiples were then added to the biofilm culture mimicking conventional TKC assessment. Timepoints measured were T0, T2, T4, T6, T8, T24 h for both planktonic (if required) and biofilm enumeration. To enumerate biofilm biomass, single glass beads were removed, placed into sterile saline to remove any planktonic cells. The bead was then transferred to a sterile bijou containing 1mL MHBII and sonicated for 30 ±5 minutes to disrupt and disperse the biofilm for enumeration. All experiments were performed in triplicate.

**Figure 3.**
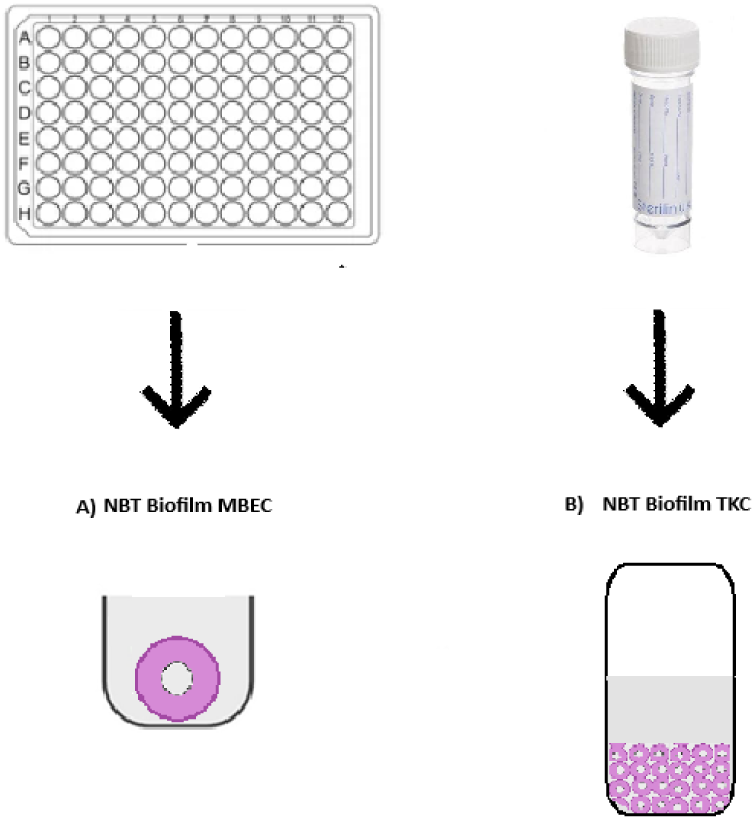
Biofilm Time Kill Curve determination.

### Measurement of antibacterial effects and pharmacodynamics

The area-under-the-bacterial-kill-curve (AUBKC log CFU/mL/h) was calculated using the log linear-trapezoidal rule for a period of 24 h (AUBKC 24), and post antibiotic exposure (after the initial 48-hour biofilm formation). The log change in viable counts compared to initial inoculum was also determined at 24 h (D24). AUBKC and D24 was related to planktonic exposures using the Sigmoid Emax Model using GraphPad Prism Version 10.3.1 (892), (San Diego, CA, USA).

## Results

Comparison of MBECs achieved using the CBD and NBT biofilm model systems for strains of *E. coli* and *S. pneumoniae* (figure 4)*, S. aureus* and *P. aeruginosa* (figure 5) and *K. pneumoniae* (figure 6) when challenged with levofloxacin, linezolid, ciprofloxacin, piperacillin-tazobactam, vancomycin and colistin, as appropriate, are shown below.

*Raw data and Mean MIC and MBECs for all strains can be found in supplementary material Table 4-6*

**Figure 4.**
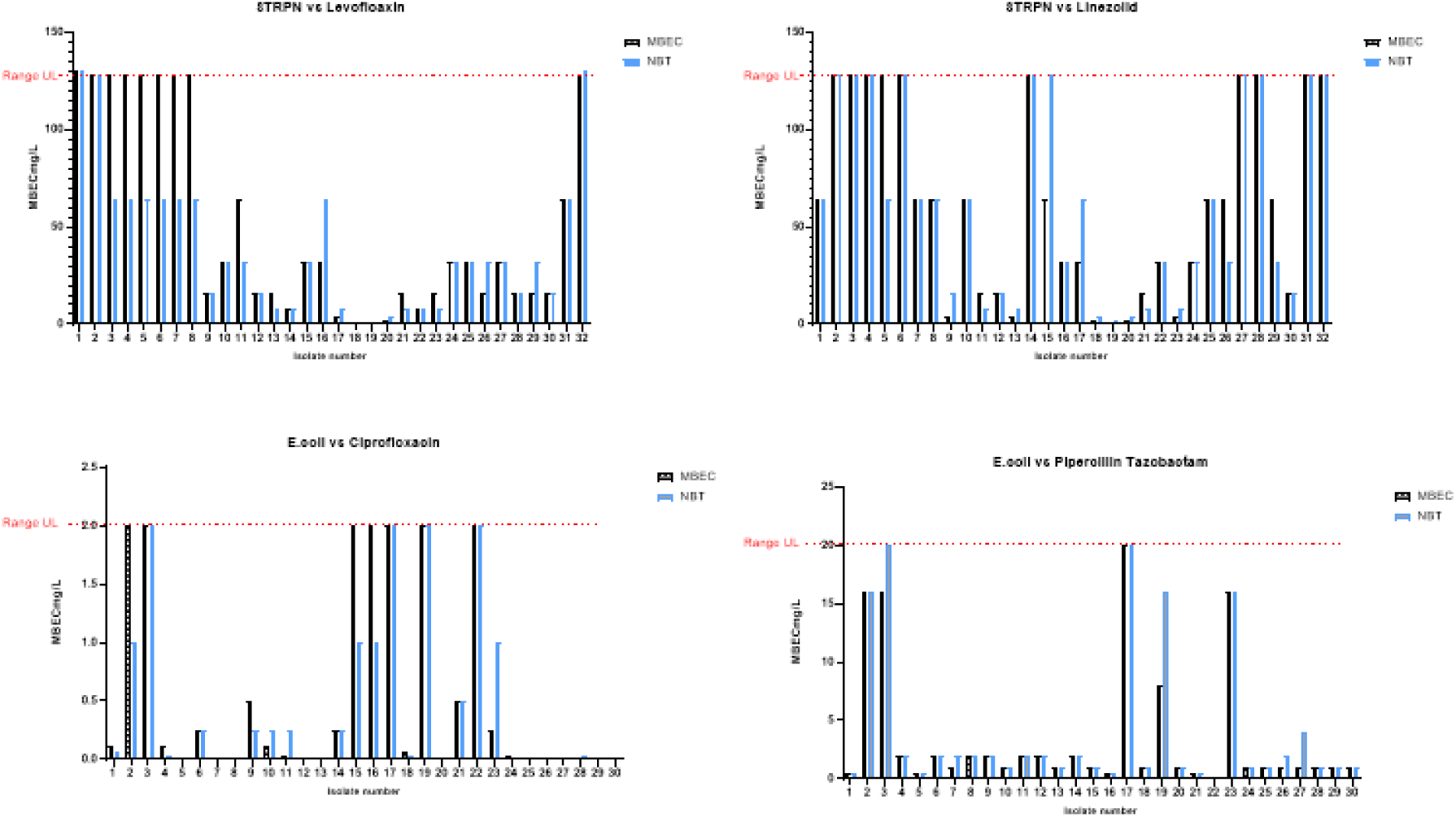
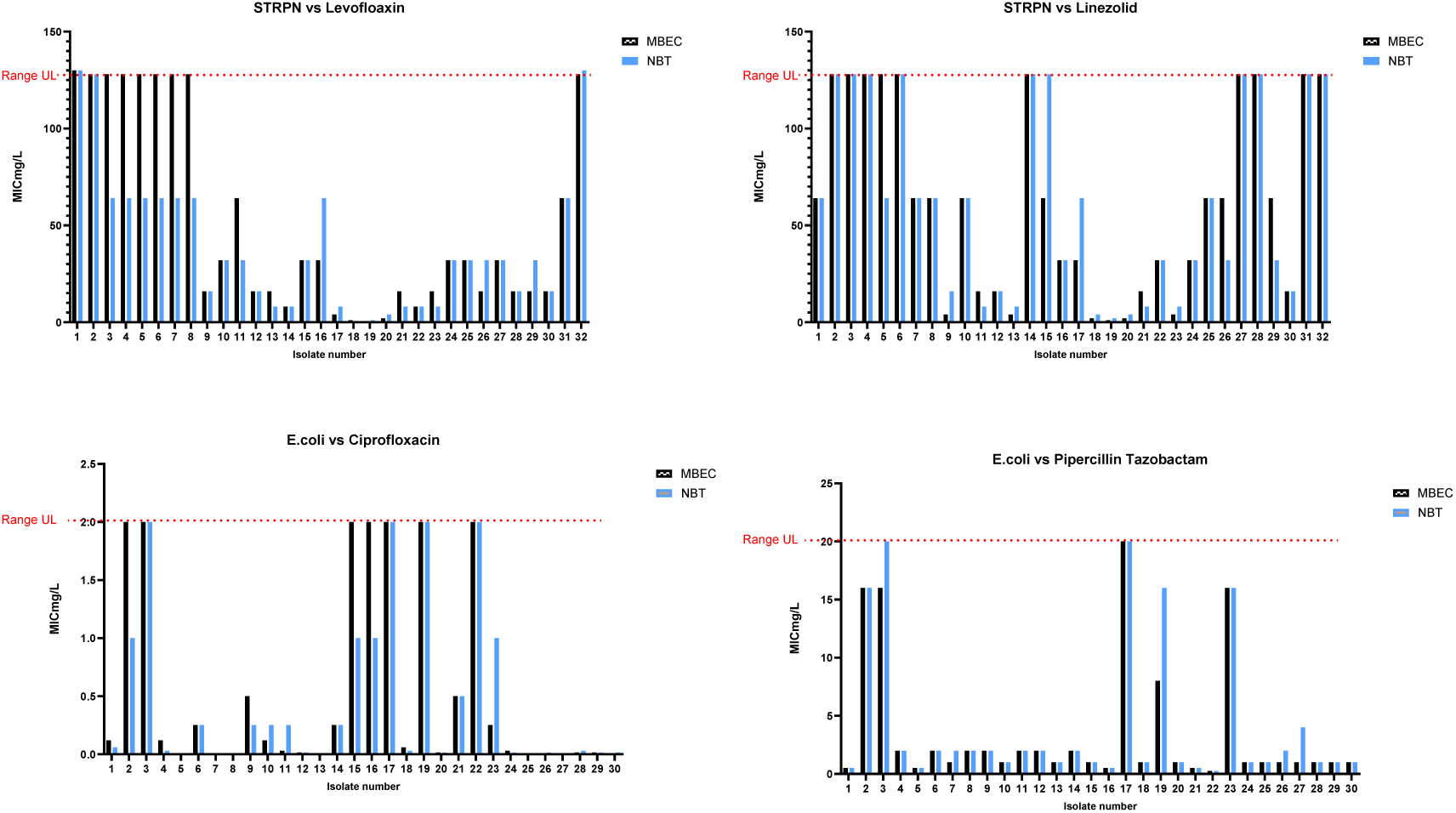
Comparison of biofilm eradication concentrations between CBD and NBT methodology S. pneumoniae *and* E. coli.

**Figure 5.**
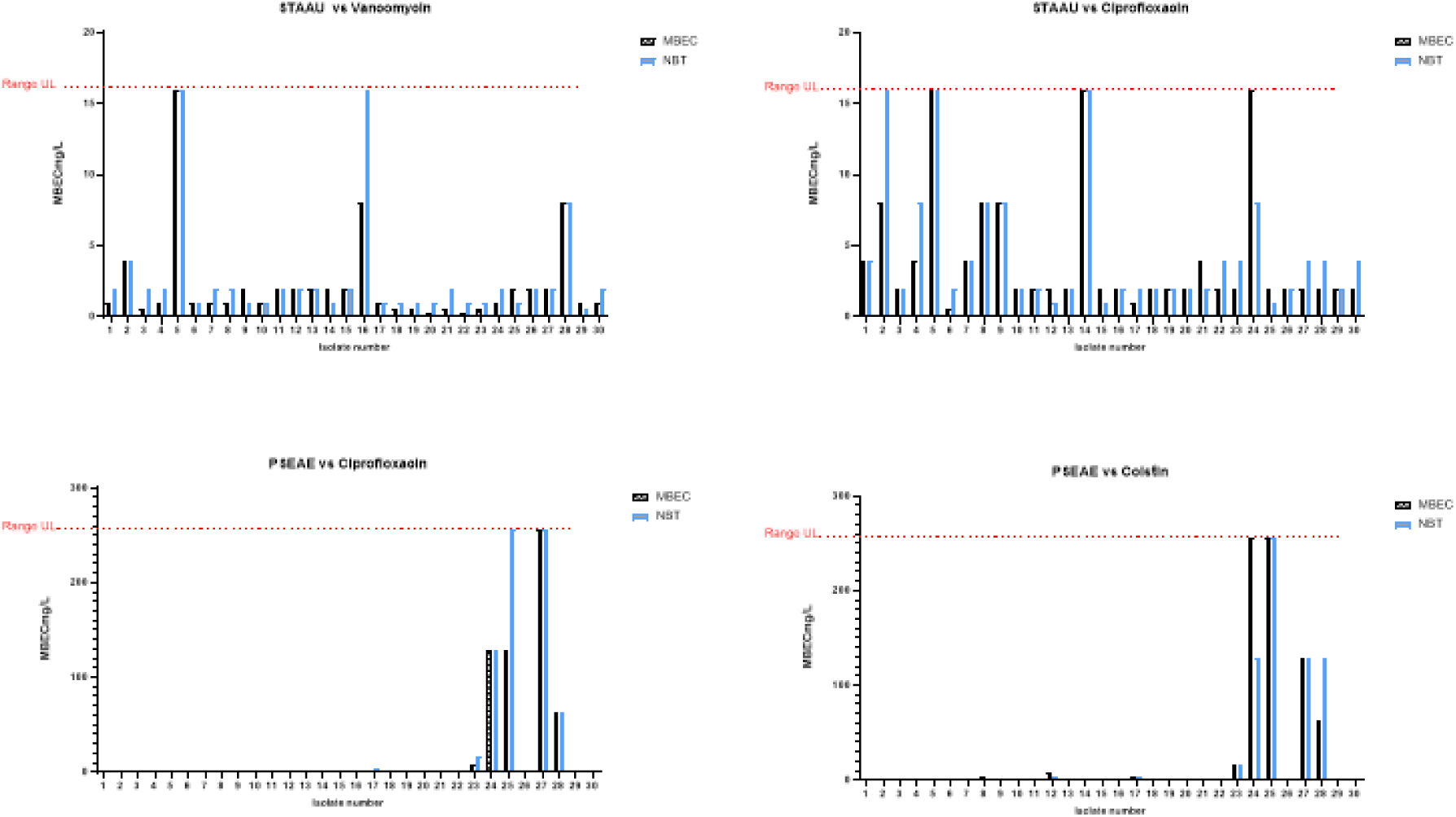
Comparison of biofilm eradication concentrations between CBD and NBT methodology *S. aureus* and *P. aeruginosa*.

**Figure 6.**
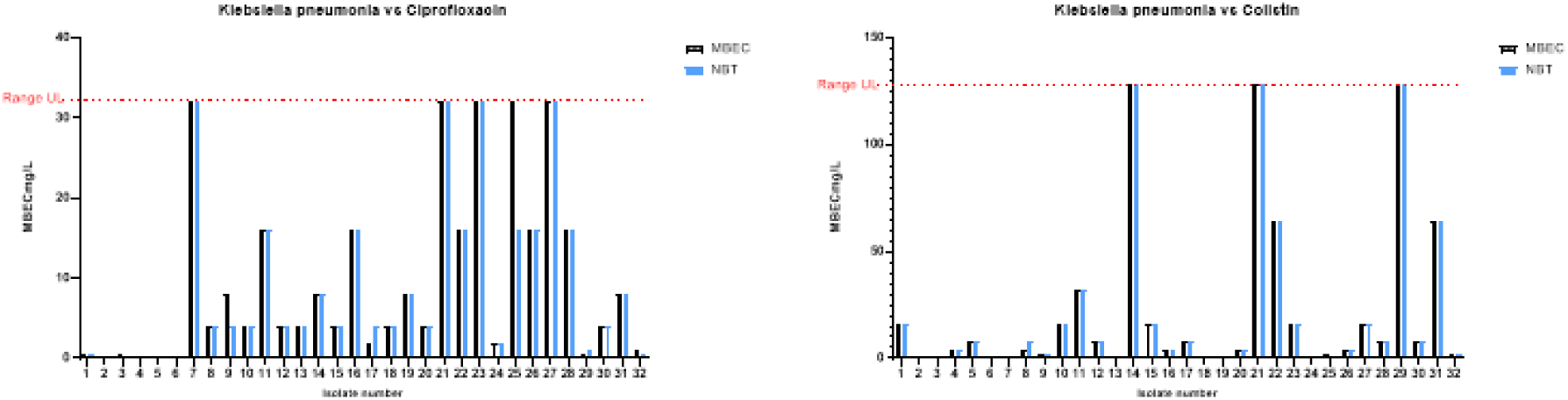
Comparison of biofilm eradication concentrations between CBD and NBT methodology *Klebsiella pneumoniae*.

The data generated here was used to inform the following TKC and BTKC assays. MIC multiples are typically used for this purpose to evaluate drug efficacy, where MIC x1 is the standard point of comparison. Higher antibiotic concentrations can assist in the determination of pharmacodynamic index (Cmax, T>MIC, AUC/MIC) and inform dosing regimens. In this application we can assess bacterial kill in both population types (planktonic and biofilm) per antibiotic concentration. Bacteria and antibiotic combinations were as followed: *K. pneumoniae* and ciprofloxacin, *S. aureus* and vancomycin and *S. pneumoniae* and levofloxacin (figure 7). All experiments were performed in triplicate for both bacterial populations.

**Figure 7.**
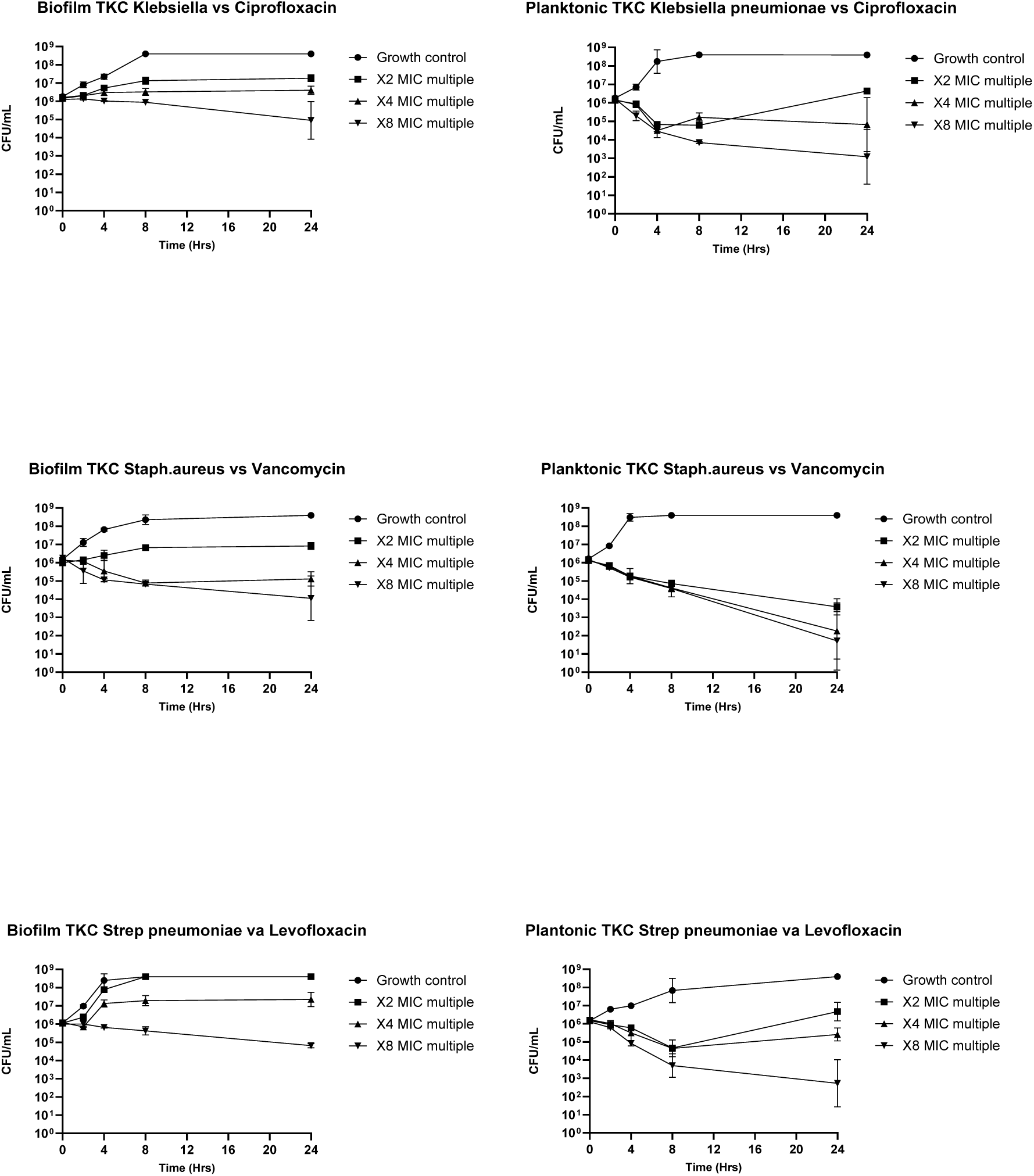
Biofilm and planktonic CFU/mL evaluations.

### Data analysis

**Table 2.** MIC and MBEC comparisons for all isolates. Values for standard MIC and NBT MBEC are presented in mg/L. Ranges correspond to differences in the determined MIC and NBT MBEC across all strains assessed. Percentage agreement was calculated as within 1 dilution factor in comparison to CBD MBEC value.

| STRPN vs Levofloxacin |  |  | STRPN vs Linezolid |  |  |
| --- | --- | --- | --- | --- | --- |
| MIC Range | 0.25-2 |  | MIC Range | 0.5-1 |  |
| MIC 50 | 1 |  | MIC 50 | 1 |  |
| MIC 90 | 2 |  | MIC 90 | 2 |  |
|  | MBEC CBD | MBEC NBT |  | MBEC CBD | MBEC NBT |
| MBEC Range | 0.5 - >128 | 0.5- > 128 | MBEC Range | 1-128 | 2-128 |
| MBEC 50 | 32 | 32 | MIC 50 | 64 | 64 |
| MBEC 90 | 128 | 64 | MIC 90 | 128 | 128 |
| % Agreement within 1 dilution | Control | 100% | % Agreement within 1 dilution | Control | 97% |
| STAAU vs Vancomycin |  |  | STAAU vs Ciprofloxacin |  |  |
| MIC Range | 0.5-1 |  | MIC Range | 0.06-0.5 |  |
| MIC 50 | 0.5 |  | MIC 50 | 0.25 |  |
| MIC 90 | 1 |  | MIC 90 | 0.5 |  |
|  | MBEC CBD | MBEC NBT |  | MBEC CBD | MBEC NBT |
| MBEC Range | 0.25- 16 | 0.5->16 | MBEC Range | 0.5-16 | 1->16 |
| MBEC 50 | 1 | 2 | MIC 50 | 2 | 2 |
| MBEC 90 | 4 | 4 | MIC 90 | 8 | 8 |
| % Agreement within 1 dilution | Control | 86% | % Agreement within 1 dilution | Control | 97% |
| ESCCO vs Ciprofloxacin |  |  | ESCCO vs Piperacillin/ Tazobactam |  |  |
| MIC Range | 0.004->1 |  | MIC Range | 0.008-8 |  |
| MIC 50 | 0.015 |  | MIC 50 | 0.06 |  |
| MIC 90 | >1 |  | MIC 90 | 4 |  |
|  | MBEC CBD | MBEC NBT |  | MBEC CBD | MBEC NBT |
| MBEC Range | 0.004->1 | 0.004->1 | MBEC Range | 0.25->16 | 0.25->16 |
| MBEC 50 | 0.06 | 0.03 | MIC 50 | 1 | 1 |
| MBEC 90 | >1 | >1 | MIC 90 | 16 | 16 |
| % Agreement within 1 dilution | Control | 90% | % Agreement within 1 dilution | Control | 97% |
| PSEAE vs Ciprofloxacin |  |  | PSEAE vs Colistin |  |  |
| MIC Range | <0.008 - 128 |  | MIC Range | 0.06-16 |  |
| MIC 50 | 0.12 |  | MIC 50 | 0.25 |  |
| MIC 90 | 8 |  | MIC 90 | 4 |  |
|  | MBEC CBD | MBEC NBT |  | MBEC CBD | MBEC NBT |
| MBEC Range | 0.06-256 | 0.06-256 | MBEC Range | 0.12-256 | 0.25-256 |
| MBEC 50 | 0.25 | 0.5 | MIC 50 | 0.5 | 1 |
| MBEC 90 | 64 | 64 | MIC 90 | 64 | 128 |
| % Agreement within 1 dilution | Control | 90% | % Agreement within 1 dilution | Control | 90% |
| **KLEPN vs Ciprofloxacin** |  |  | **KLEPN vs Colistin** |  |  |
| MIC Range | 0.008-8 |  | MIC Range | 0.25-1 |  |
| MIC 50 | 1 |  | MIC 50 | 0.25 |  |
| MIC 90 | 4 |  | MIC 90 | 0.5 |  |
|  | MBEC CBD | MBEC NBT |  | MBEC CBD | MBEC NBT |
| MBEC Range | 0.5->32 | 0.5->32 | MBEC Range | 0.5->128 | 0.5->128 |
| MBEC 50 | 4 | 4 | MIC 50 | 4 | 8 |
| MBEC 90 | >32 | >32 | MIC 90 | 64 | 64 |
| % Agreement within 1 dilution | Control | 100% | % Agreement within 1 dilution | Control | 100% |

The MIC values determined represent a distribution typical of the population of clinical isolates recovered at Southmead hospital. Biofilm sensitivity distribution is largely unknown globally. When comparing MIC 50/90 values and MBEC 50/90 values, MBEC values, typically, would be expected to be elevated in comparison due to limited antibiotic penetration, altered metabolic activity and alterations in gene expression. Observations between CBD and NBT methodology are within 1 dilution of each other for MIC 50 and 90 with exceptional % agreement (within 1 dilution) for all bacterial strains tested.

**Table 3.** Wilcoxon signed rank test all MBEC data points.

|  |  |
| --- | --- |
| Table Analysed | All data analysis |
| Column B | Data Set-B |
| vs. | vs. |
| Column A | Data Set-A |
| Wilcoxon matched-pairs signed rank test |  |
| P value | 0.0612 |
| Exact or approximate P value? | Exact |
| P value summary | ns |
| Significantly different (P < 0.05)? | No |
| One- or two-tailed P value? | Two-tailed |
| Sum of positive, negative ranks | 4914 , -3342 |
| Sum of signed ranks (W) | 1572 |
| Number of pairs | 308 |
| Number of ties (ignored) | 180 |
| Median of differences |  |
| Median | 0 |

This Wilcoxon signed rank test was used to determine if there was a statistically significant difference between the NBT and CBD methods to determine MBEC. This is a nonparametric alternative used to evaluate the median difference between two paired sample groups. These results show no significant differences as the median difference is = 0.00 and the P value is in excess of α=0.05.

**Figure 8.**
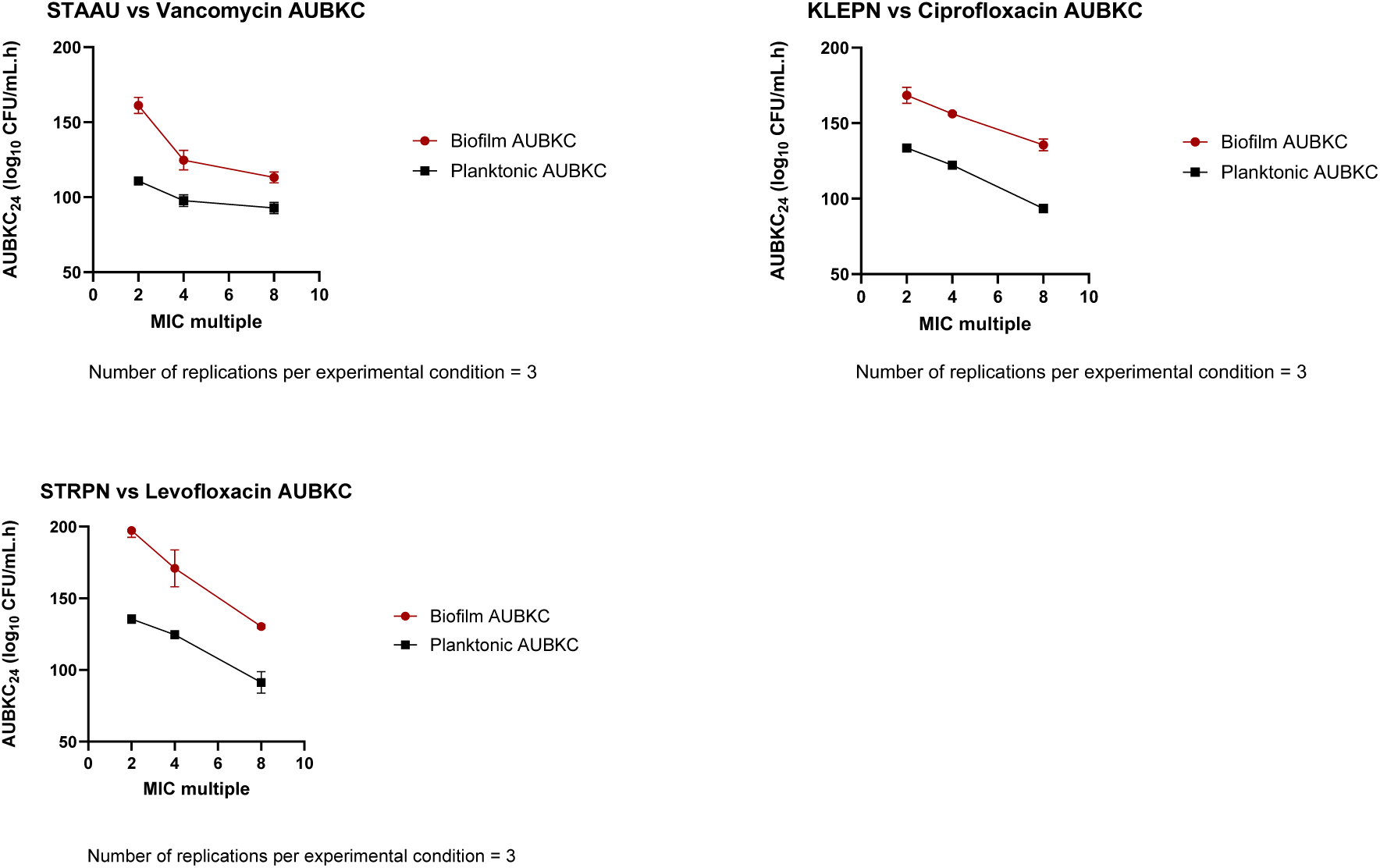
Areas under the bacterial kill curve for both biofilm and planktonic CFU enumerations at 24 hours.

The AUC was used to assess the difference in bactericidal efficacy between planktonic and biofilm cultures (Figure 8). All biofilm AUBKC were elevated in comparison to planktonic observations. Vancomycin vs STAAU specifically had superior efficacy compared to other antimicrobial-biofilm interactions. This is supported in the literature where vancomycin has enhanced ability to penetrate the biofilm and is often used in combination with other antimicrobial/non-traditional medicines to treat biofilms in prosthetic joint infections^22^ . The MBEC data correlate with data obtained by BTKC STAAU vs Vancomycin evaluations. MIC, MBEC 50/90 values and pharmacodynamic trends are all superior to other bacterial-antibiotic combinations, i.e. greater than two or more doubling dilutions and reduced bacterial population analysis.

## Conclusions

These evaluations aimed to compare two biofilm test systems (CBD MBEC® assay as the control, and NBT as the test method), and to assess biofilm correlation between static and dynamic endpoints. Whilst we acknowledge there have been other studies performed with beads in comparison to CBD method, these publications have typically focused on one specific bacterial strain^23^, anti-infective^24^ or experimental material of bead^25^. Here we describe a system developed that is appropriate for screening the activity of multiple antibacterial agents against a wide range of bacterial species in a high throughput and cost-effective manner. Observations between MIC and TKC, MBEC and BTKC endpoints show that there is an association for both types of bacteria population (Table 2 and Figure 8). Bias has been reduced where possible, for example biofilm MBEC and TKC use the same growth medium, environmental test conditions and incubation time (in line with manufacturers recommendations) for all evaluations for specific bacterial strains. This is not to say however, that other media, more reflective of specific *in vivo* environments, could not be employed in future studies, for example alteration of medium pH for gastrointestinal studies or use of synthetic mucus/sputum. The static study results revealed high levels of consistency between the two test systems for all bacterial species and drug combinations tested (within one dilutional factor compared to the commercial control), namely, *S. pneumoniae* 97%-100%, *E. coli* 90-97%, *S. aureus* 86%-97%, *P. aeruginosa* 90%, *K. pneumoniae* 100% (Table 2). We acknowledge there is subtle variation between strain and antibiotic combinations i.e. *S. aureus* vs vancomycin 86% agreement. This could be due to strain specific factors; the development of persister populations^26^, *S. aureus* biofilms entering a low metabolic state (non-uniformly) which increases the tolerance to antibtioics^27^ or the penetration of vancomycin into the biofilm, of which there is agreement with inconsistent penetration of biofilm shown in the TKC evaluations^28^.

We also acknowledge that despite generating specific bacterial species panels, indicative of biofilm formation such as original clinical antibiograms (failure of antibiotic treatment and or recurrence) and patient history (such as persisting infection > 7 days), these factors were not necessarily translatable as a predictive measure. This can be seen in some cases where MBECs were identical or within 1 dilution of MIC results per strain. The reproducibility, however, is still consistent when observing the two different methodologies. In combinations such as *E. coli* vs Piperacillin/tazobactam and *S. pneumoniae* vs Linezolid we see clear evidence of biofilm formation prior to and after the addition of drug. This appears to result in more consistency across the two test systems; 97% and 97 %, respectively.

There is no doubt that the CBD and derivations of it are enhancing the rapid screening of biofilms for various applications, yet several limitations still remain which hinder their broader adoption. Crucially, these systems lack flexibility and have no clear pathway for adaptation into more complex pharmacokinetic and pharmacodynamic evaluations. Additionally, the overall cost of single-use test systems, or the initial expense of 3D printing of similar devices, could restrict access and potentially reduce the prevalence of biofilm research within institutions.

NBT assays stand out from other biofilm systems due to the ability to generate robust data whilst offering versatile material selection, at relatively low cost. Using this method for progression of compounds through the drug development pathway is advantageous because it ensures consistency of biological techniques across static and dynamic experiments, thereby providing high-quality translational data for more complex pharmacokinetic and pharmacodynamic simulations. Importantly, the NBT assays allow global access as no specialised materials or equipment are required, making these techniques easily adoptable in resource-limited laboratory settings.

The interaction between planktonic and biofilm communities when exposed to anti-infectives is complex. Therefore, more extensive and critical observations are required to inform dosing regimens. The use of a larger culture vessel (as opposed to 96 well trays) reduces impact of nutrient depletion, allows simultaneous CFU/mL observations, (incorporates for gradients of oxygen depletion,) and assessment of rapid rate of kill and observations of bacterial regrowth which often is nuanced. Bacterial regrowth in 96 well trays, due to small culture volumes restricts nutrients, is hindered by toxins (via waste metabolite build up) and can be affected by total depletion of oxygen, which can significantly alter data analysis. Optical density evaluations which are commonly employed with planktonic TKCs cannot be applied for biofilm bacterial population assessment due to EPS, sediment interfering with light absorption (or scattering) and the inability to differentiate planktonic cells and biofilm cells^29^. Therefore, the use of larger culture vessels, as described in this manuscript, can generate enhanced data sets which could be used to provide an intermediate screening method and can provide indications of pharmacodynamic drivers such as C_max_, T>MIC or AUC dependant killing^30^. Whilst planktonic pharmacodynamics are widely established for a majority of commonly used antimicrobials, biofilm drivers remain less defined^31^. These techniques could begin to bridge this gap to ensure optimum dosing regimens with a reduced risk of treatment failure or relapse.

Such systems could include a wider variety of single and polymicrobial biofilm infections. This framework can also extend to novel therapeutic combinations, including bacteriophage and antibiotic-phage synergy where pharmacodynamics is perhaps more valuable as an indicator for therapeutic success in comparison to static tests.

The complexity of diagnosing and treating biofilm-associated infections necessitates a comprehensive strategy to mitigate the rising rates of AMR. The methods presented here offer a robust, reproducible alternative and include adaptable systems which can be implemented globally at low cost and use standard laboratory equipment providing fuller data sets which are translatable to clinical scenarios.

---

**Supplementary Table 1.**
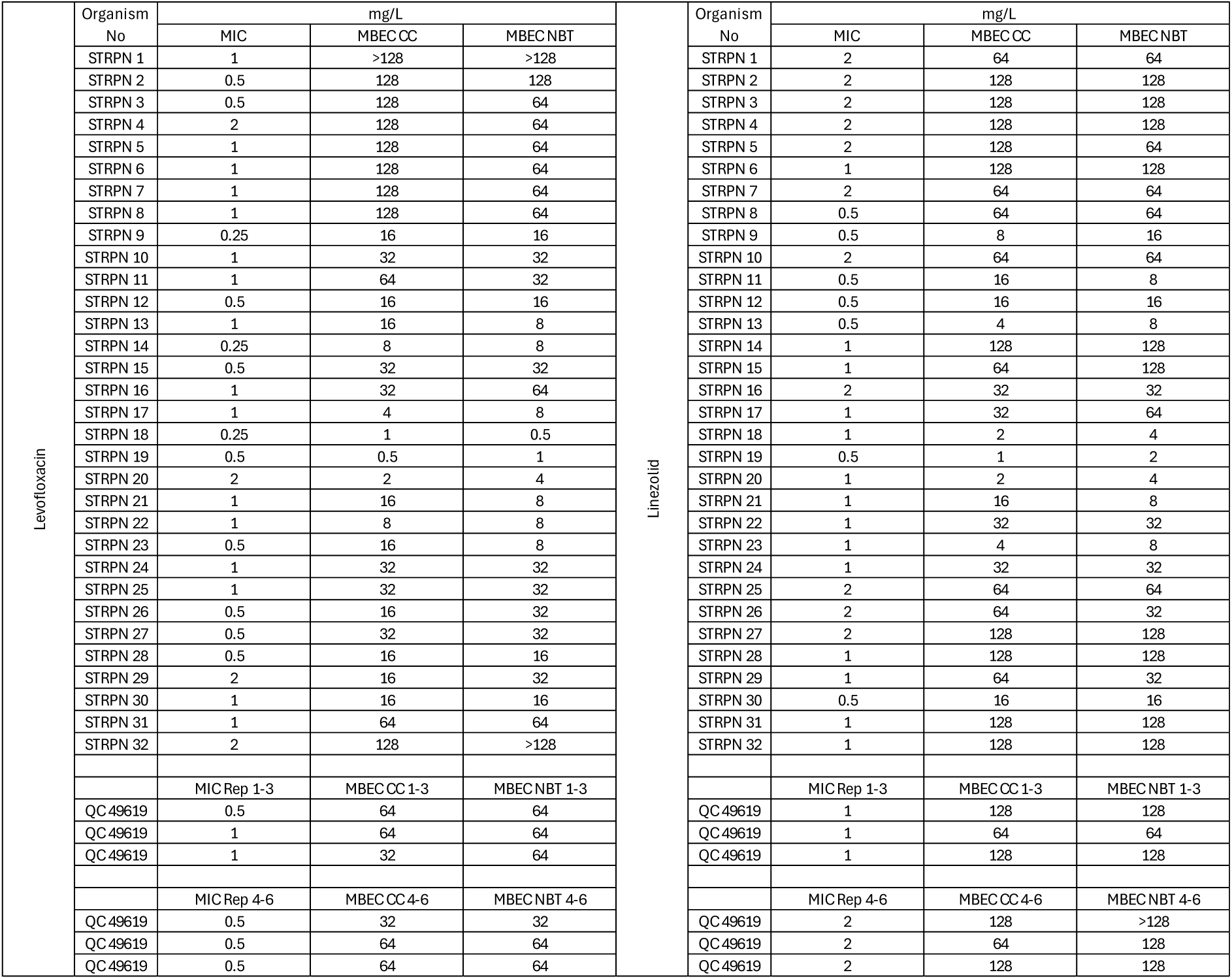
Mean MIC and MBEC STRPN vs Levofloxacin, Mean MIC and MBEC STRPN vs Linezolid.

**Supplementary Table 2.**
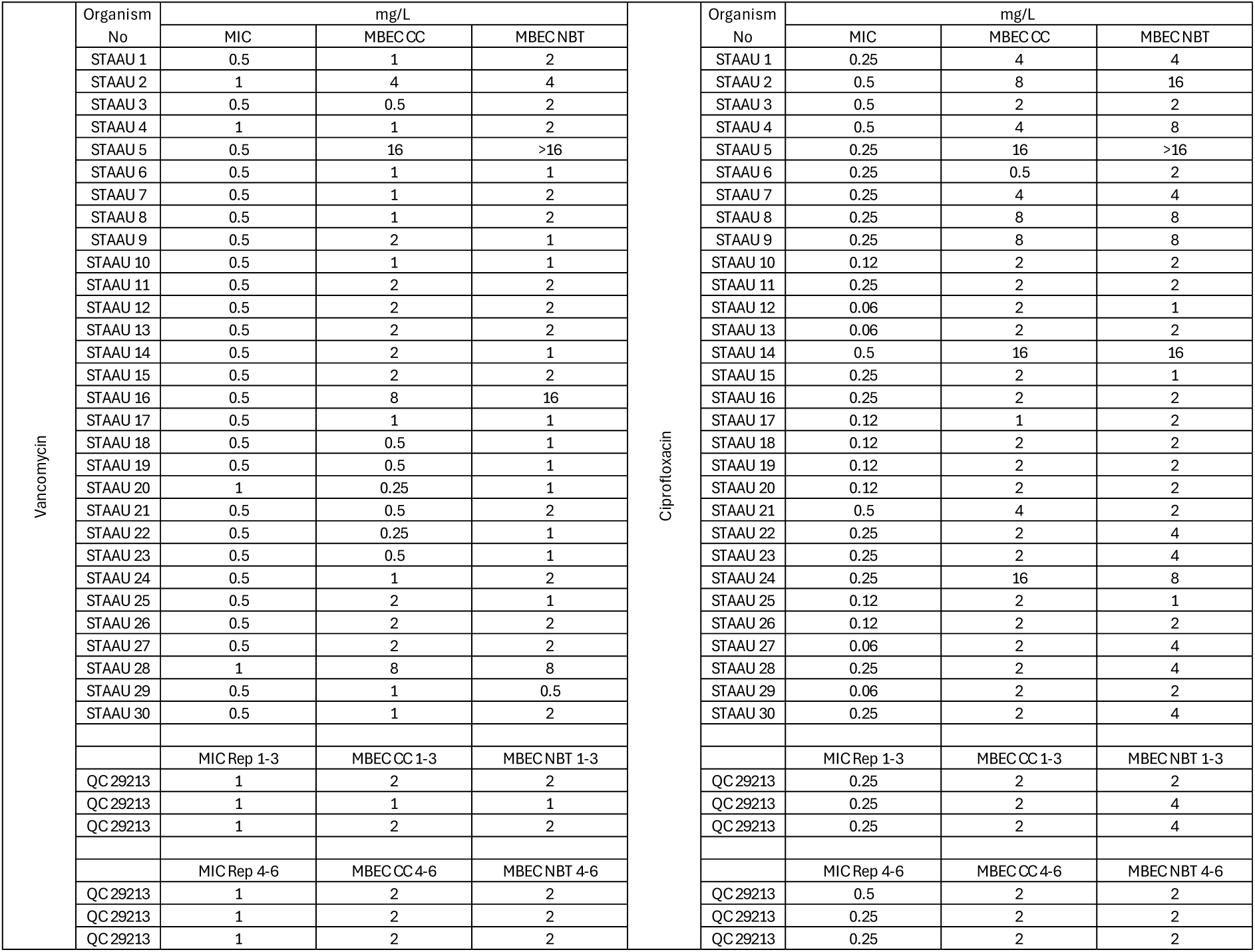
Mean MIC and MBEC Staph. aureus vs Vancomycin, Mean MIC and MBEC Staph. Aureus vs ciprofloxacin.

|  | Organism No | mg/L |  |  |  | Organism No | mg/L |  |  |
| --- | --- | --- | --- | --- | --- | --- | --- | --- | --- |
|  |  | MIC | MBECCC | MBECNBT |  |  | MIC | MBECCC | MBECNBT |
| Vancomycin | STAAU 1 | 0.5 | 1 | 2 | Ciprofloxacin | STAAU 1 | 0.25 | 4 | 4 |
|  | STAAU 2 | 1 | 4 | 4 |  | STAAU 2 | 0.5 | 8 | 16 |
|  | STAAU 3 | 0.5 | 0.5 | 2 |  | STAAU 3 | 0.5 | 2 | 2 |
|  | STAAU 4 | 1 | 1 | 2 |  | STAAU 4 | 0.5 | 4 | 8 |
|  | STAAU 5 | 0.5 | 16 | >16 |  | STAAU 5 | 0.25 | 16 | >16 |
|  | STAAU 6 | 0.5 | 1 | 1 |  | STAAU 6 | 0.25 | 0.5 | 2 |
|  | STAAU 7 | 0.5 | 1 | 2 |  | STAAU 7 | 0.25 | 4 | 4 |
|  | STAAU 8 | 0.5 | 1 | 2 |  | STAAU 8 | 0.25 | 8 | 8 |
|  | STAAU 9 | 0.5 | 2 | 1 |  | STAAU 9 | 0.25 | 8 | 8 |
|  | STAAU 10 | 0.5 | 1 | 1 |  | STAAU 10 | 0.12 | 2 | 2 |
|  | STAAU 11 | 0.5 | 2 | 2 |  | STAAU 11 | 0.25 | 2 | 2 |
|  | STAAU 12 | 0.5 | 2 | 2 |  | STAAU 12 | 0.06 | 2 | 1 |
|  | STAAU 13 | 0.5 | 2 | 2 |  | STAAU 13 | 0.06 | 2 | 2 |
|  | STAAU 14 | 0.5 | 2 | 1 |  | STAAU 14 | 0.5 | 16 | 16 |
|  | STAAU 15 | 0.5 | 2 | 2 |  | STAAU 15 | 0.25 | 2 | 1 |
|  | STAAU 16 | 0.5 | 8 | 16 |  | STAAU 16 | 0.25 | 2 | 2 |
|  | STAAU 17 | 0.5 | 1 | 1 |  | STAAU 17 | 0.12 | 1 | 2 |
|  | STAAU 18 | 0.5 | 0.5 | 1 |  | STAAU 18 | 0.12 | 2 | 2 |
|  | STAAU 19 | 0.5 | 0.5 | 1 |  | STAAU 19 | 0.12 | 2 | 2 |
|  | STAAU 20 | 1 | 0.25 | 1 |  | STAAU 20 | 0.12 | 2 | 2 |
|  | STAAU 21 | 0.5 | 0.5 | 2 |  | STAAU 21 | 0.5 | 4 | 2 |
|  | STAAU 22 | 0.5 | 0.25 | 1 |  | STAAU 22 | 0.25 | 2 | 4 |
|  | STAAU 23 | 0.5 | 0.5 | 1 |  | STAAU 23 | 0.25 | 2 | 4 |
|  | STAAU 24 | 0.5 | 1 | 2 |  | STAAU 24 | 0.25 | 16 | 8 |
|  | STAAU 25 | 0.5 | 2 | 1 |  | STAAU 25 | 0.12 | 2 | 1 |
|  | STAAU 26 | 0.5 | 2 | 2 |  | STAAU 26 | 0.12 | 2 | 2 |
|  | STAAU 27 | 0.5 | 2 | 2 |  | STAAU 27 | 0.06 | 2 | 4 |
|  | STAAU 28 | 1 | 8 | 8 |  | STAAU 28 | 0.25 | 2 | 4 |
|  | STAAU 29 | 0.5 | 1 | 0.5 |  | STAAU 29 | 0.06 | 2 | 2 |
|  | STAAU 30 | 0.5 | 1 | 2 |  | STAAU 30 | 0.25 | 2 | 4 |
|  |  | MIC Rep 1-3 | MBECCC 1-3 | MBECNBT 1-3 |  |  | MIC Rep 1-3 | MBECCC 1-3 | MBECNBT 1-3 |
|  | QC 29213 | 1 | 2 | 2 |  | QC 29213 | 0.25 | 2 | 2 |
|  | QC 29213 | 1 | 1 | 1 |  | QC 29213 | 0.25 | 2 | 4 |
|  | QC 29213 | 1 | 2 | 2 |  | QC 29213 | 0.25 | 2 | 4 |
|  |  | MIC Rep 4-6 | MBECCC 4-6 | MBECNBT 4-6 |  |  | MIC Rep 4-6 | MBECCC 4-6 | MBECNBT 4-6 |
|  | QC 29213 | 1 | 2 | 2 |  | QC 29213 | 0.5 | 2 | 2 |
|  | QC 29213 | 1 | 2 | 2 |  | QC 29213 | 0.25 | 2 | 2 |
|  | QC 29213 | 1 | 2 | 2 |  | QC 29213 | 0.25 | 2 | 2 |

**Supplementary Table 3.**
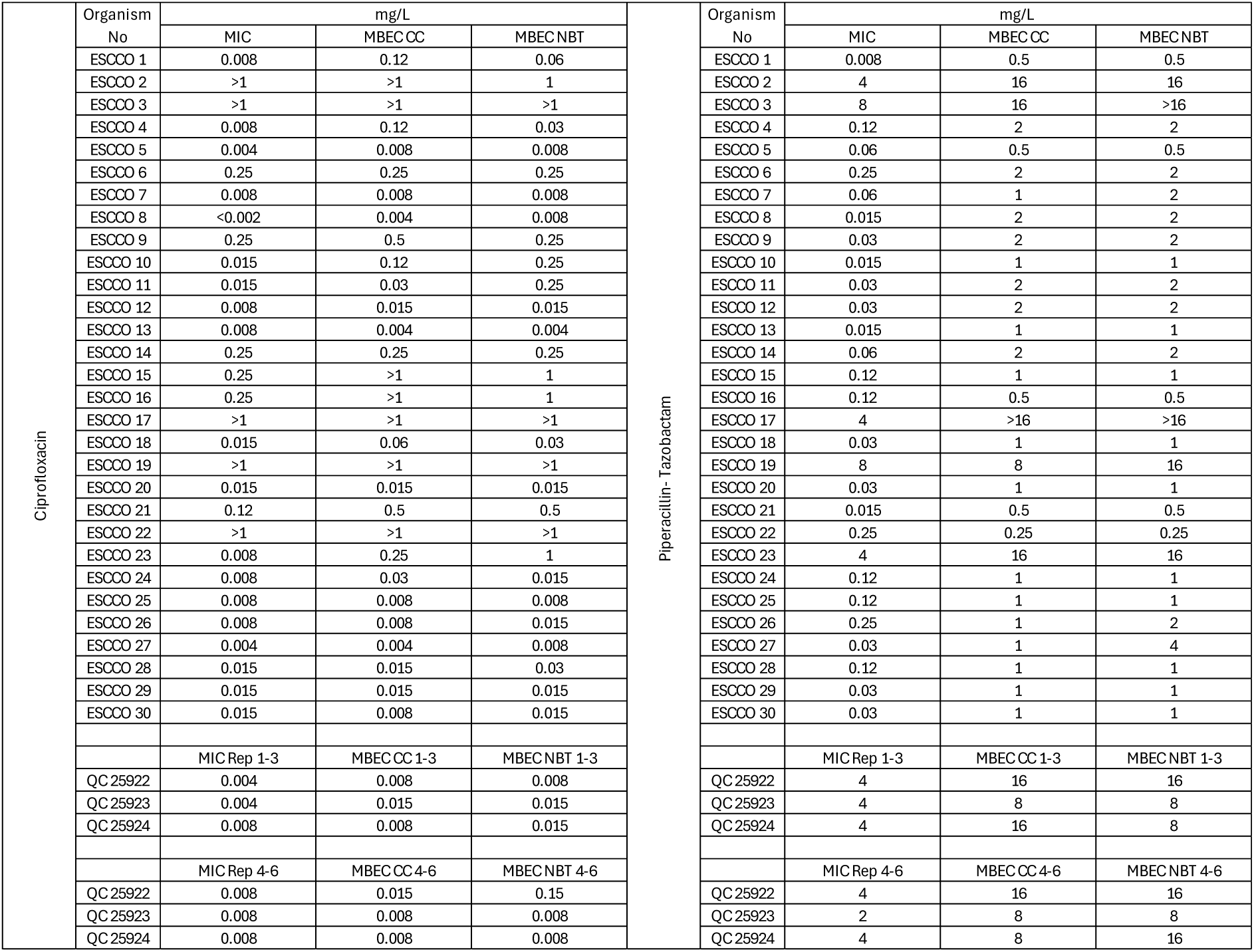
Mean MIC and MBEC E. coli vs Ciprofloxacin, Mean MIC and MBEC E.coli vs Piperacillin/Tazobactam.

**Supplementary Table 4.**
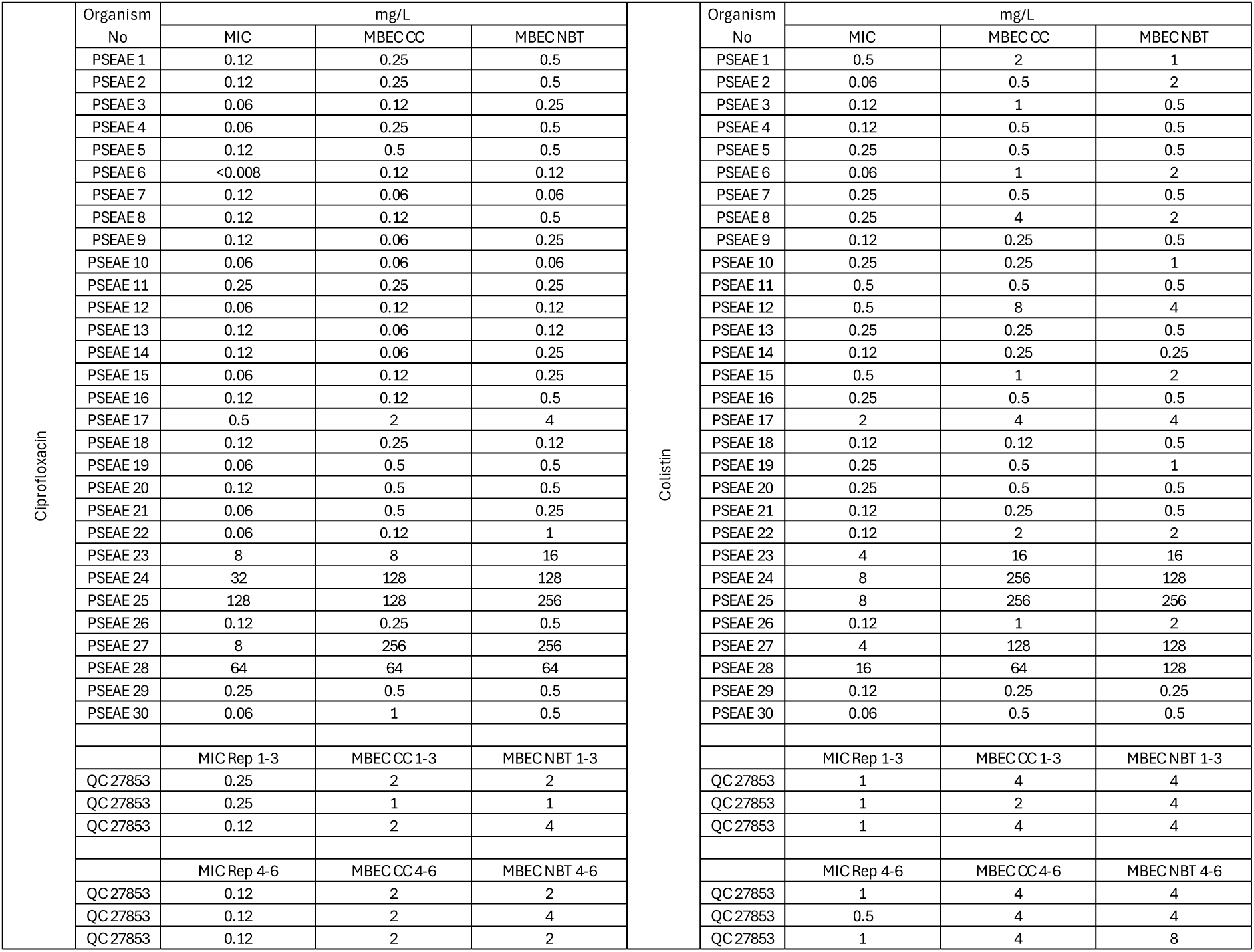
Mean MIC and MBEC Pseudomonas aeruginosa vs ciprofloxacin, Mean MIC MBEC Pseudomonas aeruginosa vs Colistin.

**Supplementary Table 5.**
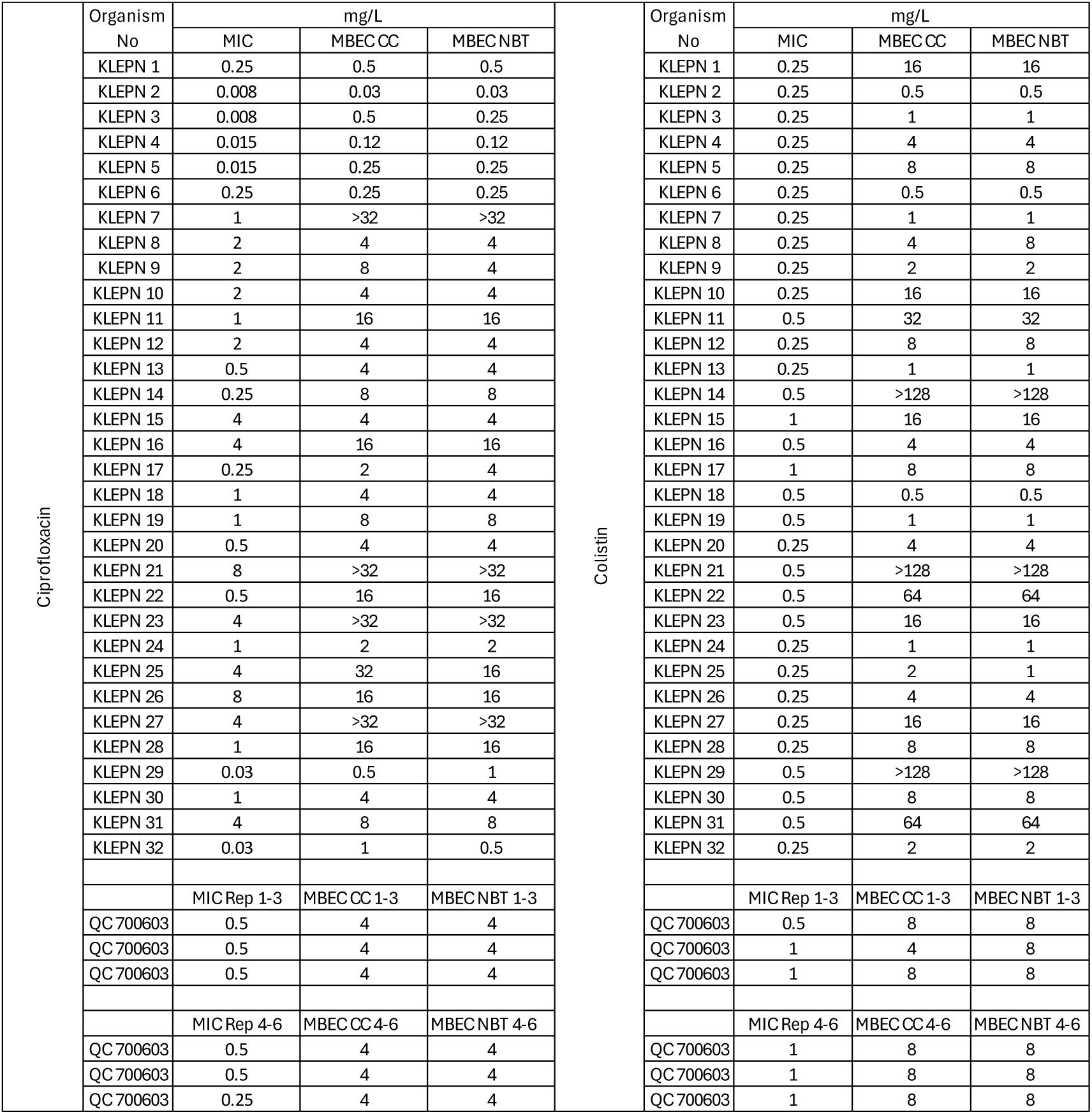
Mean MIC and MBEC Klebsiella pneumonia vs ciprofloxacin, Mean MIC and MBEC Klebsiella pneumonia vs Colistin.

|  | Organism No | mg/L |  |  |
| --- | --- | --- | --- | --- |
|  |  | MIC | MBECCC | MBECNBT |
| Ciprofloxacin | KLEPN 1 | 0.25 | 0.5 | 0.5 |
|  | KLEPN 2 | 0.008 | 0.03 | 0.03 |
|  | KLEPN 3 | 0.008 | 0.5 | 0.25 |
|  | KLEPN 4 | 0.015 | 0.12 | 0.12 |
|  | KLEPN 5 | 0.015 | 0.25 | 0.25 |
|  | KLEPN 6 | 0.25 | 0.25 | 0.25 |
|  | KLEPN 7 | 1 | >32 | >32 |
|  | KLEPN 8 | 2 | 4 | 4 |
|  | KLEPN 9 | 2 | 8 | 4 |
|  | KLEPN 10 | 2 | 4 | 4 |
|  | KLEPN 11 | 1 | 16 | 16 |
|  | KLEPN 12 | 2 | 4 | 4 |
|  | KLEPN 13 | 0.5 | 4 | 4 |
|  | KLEPN 14 | 0.25 | 8 | 8 |
|  | KLEPN 15 | 4 | 4 | 4 |
|  | KLEPN 16 | 4 | 16 | 16 |
|  | KLEPN 17 | 0.25 | 2 | 4 |
|  | KLEPN 18 | 1 | 4 | 4 |
|  | KLEPN 19 | 1 | 8 | 8 |
|  | KLEPN 20 | 0.5 | 4 | 4 |
|  | KLEPN 21 | 8 | >32 | >32 |
|  | KLEPN 22 | 0.5 | 16 | 16 |
|  | KLEPN 23 | 4 | >32 | >32 |
|  | KLEPN 24 | 1 | 2 | 2 |
|  | KLEPN 25 | 4 | 32 | 16 |
|  | KLEPN 26 | 8 | 16 | 16 |
|  | KLEPN 27 | 4 | >32 | >32 |
|  | KLEPN 28 | 1 | 16 | 16 |
|  | KLEPN 29 | 0.03 | 0.5 | 1 |
|  | KLEPN 30 | 1 | 4 | 4 |
|  | KLEPN 31 | 4 | 8 | 8 |
|  | KLEPN 32 | 0.03 | 1 | 0.5 |
|  |  | MIC Rep 1-3 | MBECCC 1-3 | MBECNBT 1-3 |
|  | QC 700603 | 0.5 | 4 | 4 |
|  | QC 700603 | 0.5 | 4 | 4 |
|  | QC 700603 | 0.5 | 4 | 4 |
|  |  | MIC Rep 4-6 | MBECCC 4-6 | MBECNBT 4-6 |
|  | QC 700603 | 0.5 | 4 | 4 |
|  | QC 700603 | 0.5 | 4 | 4 |
|  | QC 700603 | 0.25 | 4 | 4 |
| Colistin | Organism No | mg/L |  |  |
|  |  | MIC | MBECCC | MBECNBT |
|  | KLEPN 1 | 0.25 | 16 | 16 |
|  | KLEPN 2 | 0.25 | 0.5 | 0.5 |
|  | KLEPN 3 | 0.25 | 1 | 1 |
|  | KLEPN 4 | 0.25 | 4 | 4 |
|  | KLEPN 5 | 0.25 | 8 | 8 |
|  | KLEPN 6 | 0.25 | 0.5 | 0.5 |
|  | KLEPN 7 | 0.25 | 1 | 1 |
|  | KLEPN 8 | 0.25 | 4 | 8 |
|  | KLEPN 9 | 0.25 | 2 | 2 |
|  | KLEPN 10 | 0.25 | 16 | 16 |
|  | KLEPN 11 | 0.5 | 32 | 32 |
|  | KLEPN 12 | 0.25 | 8 | 8 |
|  | KLEPN 13 | 0.25 | 1 | 1 |
|  | KLEPN 14 | 0.5 | >128 | >128 |
|  | KLEPN 15 | 1 | 16 | 16 |
|  | KLEPN 16 | 0.5 | 4 | 4 |
|  | KLEPN 17 | 1 | 8 | 8 |
|  | KLEPN 18 | 0.5 | 0.5 | 0.5 |
|  | KLEPN 19 | 0.5 | 1 | 1 |
|  | KLEPN 20 | 0.25 | 4 | 4 |
|  | KLEPN 21 | 0.5 | >128 | >128 |
|  | KLEPN 22 | 0.5 | 64 | 64 |
|  | KLEPN 23 | 0.5 | 16 | 16 |
|  | KLEPN 24 | 0.25 | 1 | 1 |
|  | KLEPN 25 | 0.25 | 2 | 1 |
|  | KLEPN 26 | 0.25 | 4 | 4 |
|  | KLEPN 27 | 0.25 | 16 | 16 |
|  | KLEPN 28 | 0.25 | 8 | 8 |
|  | KLEPN 29 | 0.5 | >128 | >128 |
|  | KLEPN 30 | 0.5 | 8 | 8 |
|  | KLEPN 31 | 0.5 | 64 | 64 |
|  | KLEPN 32 | 0.25 | 2 | 2 |
|  |  | MIC Rep 1-3 | MBECCC 1-3 | MBECNBT 1-3 |
|  | QC 700603 | 0.5 | 8 | 8 |
|  | QC 700603 | 1 | 4 | 8 |
|  | QC 700603 | 1 | 8 | 8 |
|  |  | MIC Rep 4-6 | MBECCC 4-6 | MBECNBT 4-6 |
|  | QC 700603 | 1 | 8 | 8 |
|  | QC 700603 | 1 | 8 | 8 |
|  | QC 700603 | 1 | 8 | 8 |

**Sup table 6.**
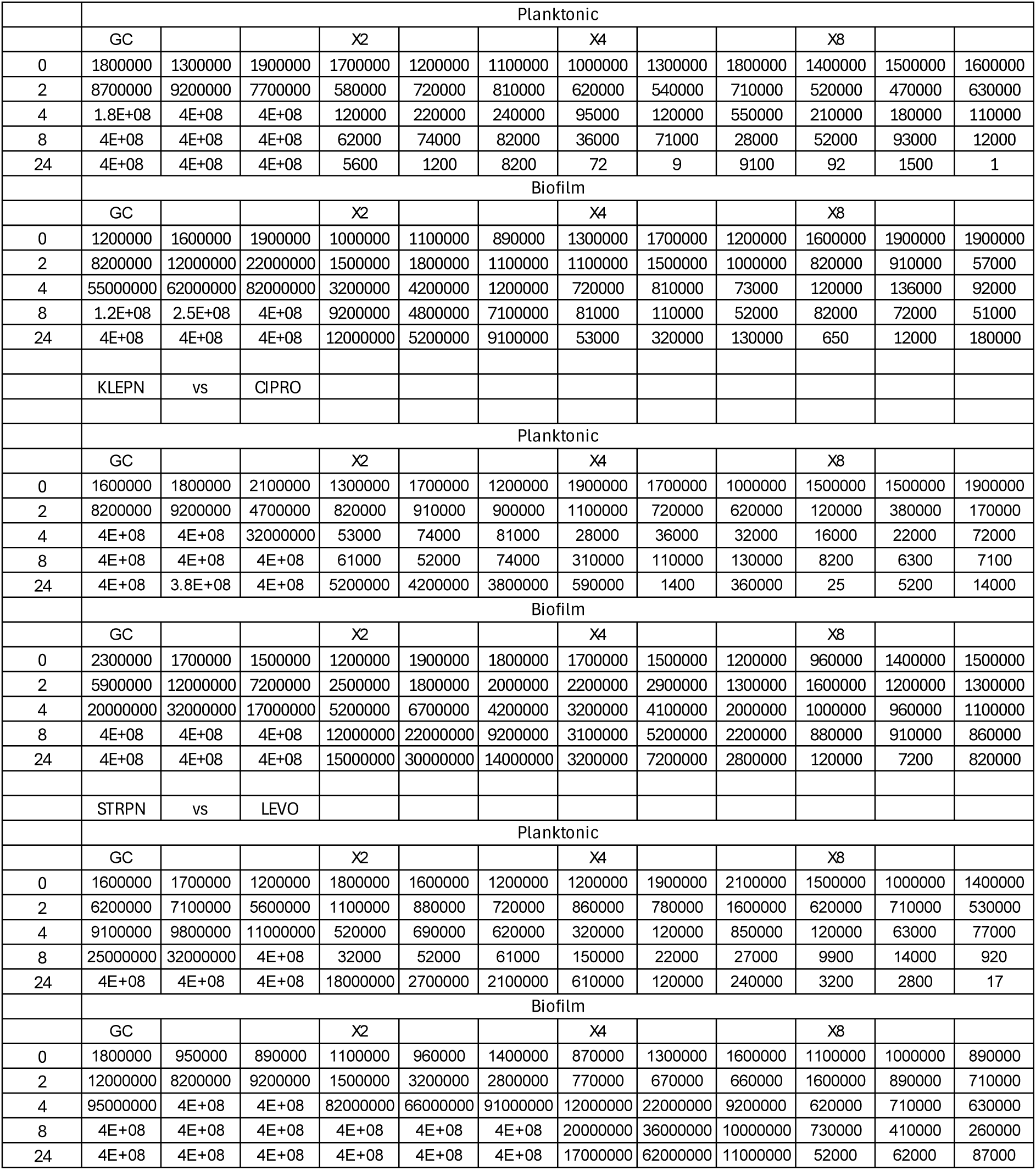
CFU/mL raw data.

**Supplementary figure 1.**
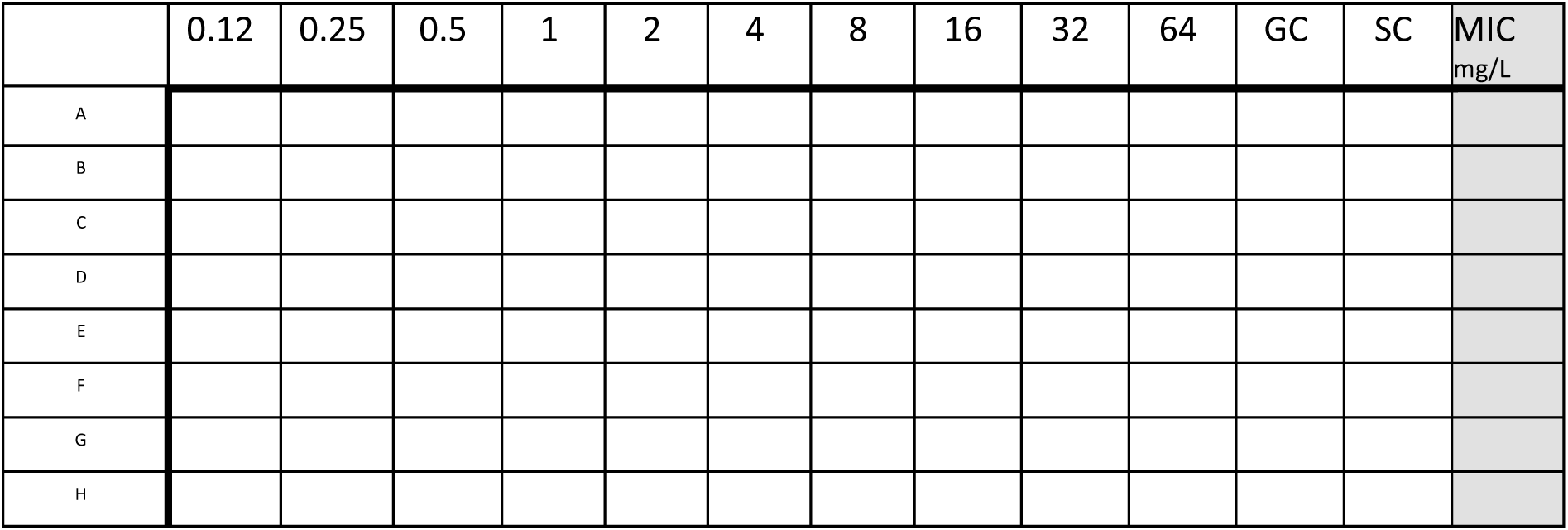
Typical challenge plate

