## Supplemental tables for "Development of low cost, robust and reproducible biofilm static and pharmacodynamic assays"

Supplementary Table 1 – Mean MIC and MBEC STRPN vs Levofloxacin , Mean MIC and MBEC STRPN vs Linezolid

**Levofloxacin**

| Organism No | mg/L MIC | mg/L MBEC CC | mg/L MBEC NBT |
|---|---|---|---|
| STRPN 1 | 1 | >128 | >128 |
| STRPN 2 | 0.5 | 128 | 128 |
| STRPN 3 | 0.5 | 128 | 64 |
| STRPN 4 | 2 | 128 | 64 |
| STRPN 5 | 1 | 128 | 64 |
| STRPN 6 | 1 | 128 | 64 |
| STRPN 7 | 1 | 128 | 64 |
| STRPN 8 | 1 | 128 | 64 |
| STRPN 9 | 0.25 | 16 | 16 |
| STRPN 10 | 1 | 32 | 32 |
| STRPN 11 | 1 | 64 | 32 |
| STRPN 12 | 0.5 | 16 | 16 |
| STRPN 13 | 1 | 16 | 8 |
| STRPN 14 | 0.25 | 8 | 8 |
| STRPN 15 | 0.5 | 32 | 32 |
| STRPN 16 | 1 | 32 | 64 |
| STRPN 17 | 1 | 4 | 8 |
| STRPN 18 | 0.25 | 1 | 0.5 |
| STRPN 19 | 0.5 | 0.5 | 1 |
| STRPN 20 | 2 | 2 | 4 |
| STRPN 21 | 1 | 16 | 8 |
| STRPN 22 | 1 | 8 | 8 |
| STRPN 23 | 0.5 | 16 | 8 |
| STRPN 24 | 1 | 32 | 32 |
| STRPN 25 | 1 | 32 | 32 |
| STRPN 26 | 0.5 | 16 | 32 |
| STRPN 27 | 0.5 | 32 | 32 |
| STRPN 28 | 0.5 | 16 | 16 |
| STRPN 29 | 2 | 16 | 32 |
| STRPN 30 | 1 | 16 | 16 |
| STRPN 31 | 1 | 64 | 64 |
| STRPN 32 | 2 | 128 | >128 |
| | MIC Rep 1-3 | MBEC CC 1-3 | MBEC NBT 1-3 |
| QC 49619 | 0.5 | 64 | 64 |
| QC 49619 | 1 | 64 | 64 |
| QC 49619 | 1 | 32 | 64 |
| | MIC Rep 4-6 | MBEC CC 4-6 | MBEC NBT 4-6 |
| QC 49619 | 0.5 | 32 | 32 |
| QC 49619 | 0.5 | 64 | 64 |
| QC 49619 | 0.5 | 64 | 64 |

**Linezolid**

| Organism No | mg/L MIC | mg/L MBEC CC | mg/L MBEC NBT |
|---|---|---|---|
| STRPN 1 | 2 | 64 | 64 |
| STRPN 2 | 2 | 128 | 128 |
| STRPN 3 | 2 | 128 | 128 |
| STRPN 4 | 2 | 128 | 128 |
| STRPN 5 | 2 | 128 | 64 |
| STRPN 6 | 1 | 128 | 128 |
| STRPN 7 | 2 | 64 | 64 |
| STRPN 8 | 0.5 | 64 | 64 |
| STRPN 9 | 0.5 | 8 | 16 |
| STRPN 10 | 2 | 64 | 64 |
| STRPN 11 | 0.5 | 16 | 8 |
| STRPN 12 | 0.5 | 16 | 16 |
| STRPN 13 | 0.5 | 4 | 8 |
| STRPN 14 | 1 | 128 | 128 |
| STRPN 15 | 1 | 64 | 128 |
| STRPN 16 | 2 | 32 | 32 |
| STRPN 17 | 1 | 32 | 64 |
| STRPN 18 | 1 | 2 | 4 |
| STRPN 19 | 0.5 | 1 | 2 |
| STRPN 20 | 1 | 2 | 4 |
| STRPN 21 | 1 | 16 | 8 |
| STRPN 22 | 1 | 32 | 32 |
| STRPN 23 | 1 | 4 | 8 |
| STRPN 24 | 1 | 32 | 32 |
| STRPN 25 | 2 | 64 | 64 |
| STRPN 26 | 2 | 64 | 32 |
| STRPN 27 | 2 | 128 | 128 |
| STRPN 28 | 1 | 128 | 128 |
| STRPN 29 | 1 | 64 | 32 |
| STRPN 30 | 0.5 | 16 | 16 |
| STRPN 31 | 1 | 128 | 128 |
| STRPN 32 | 1 | 128 | 128 |
| | MIC Rep 1-3 | MBEC CC 1-3 | MBEC NBT 1-3 |
| QC 49619 | 1 | 128 | 128 |
| QC 49619 | 1 | 64 | 64 |
| QC 49619 | 1 | 128 | 128 |
| | MIC Rep 4-6 | MBEC CC 4-6 | MBEC NBT 4-6 |
| QC 49619 | 2 | 128 | >128 |
| QC 49619 | 2 | 64 | 128 |
| QC 49619 | 2 | 128 | 128 |

Supplementary Table 2 – Mean MIC and MBEC Staph. aureus vs Vancomycin , Mean MIC and MBEC Staph. Aureus vs ciprofloxacin

| | Organism No | mg/L | | | | Organism No | mg/L | | |
|---|---|---|---|---|---|---|---|---|---|
| | | MIC | MBEC CC | MBEC NBT | | | MIC | MBEC CC | MBEC NBT |
| Vancomycin | STAAU 1 | 0.5 | 1 | 2 | Ciprofloxacin | STAAU 1 | 0.25 | 4 | 4 |
| | STAAU 2 | 1 | 4 | 4 | | STAAU 2 | 0.5 | 8 | 16 |
| | STAAU 3 | 0.5 | 0.5 | 2 | | STAAU 3 | 0.5 | 2 | 2 |
| | STAAU 4 | 1 | 1 | 2 | | STAAU 4 | 0.5 | 4 | 8 |
| | STAAU 5 | 0.5 | 16 | >16 | | STAAU 5 | 0.25 | 16 | >16 |
| | STAAU 6 | 0.5 | 1 | 1 | | STAAU 6 | 0.25 | 0.5 | 2 |
| | STAAU 7 | 0.5 | 1 | 2 | | STAAU 7 | 0.25 | 4 | 4 |
| | STAAU 8 | 0.5 | 1 | 2 | | STAAU 8 | 0.25 | 8 | 8 |
| | STAAU 9 | 0.5 | 2 | 1 | | STAAU 9 | 0.25 | 8 | 8 |
| | STAAU 10 | 0.5 | 1 | 1 | | STAAU 10 | 0.12 | 2 | 2 |
| | STAAU 11 | 0.5 | 2 | 2 | | STAAU 11 | 0.25 | 2 | 2 |
| | STAAU 12 | 0.5 | 2 | 2 | | STAAU 12 | 0.06 | 2 | 1 |
| | STAAU 13 | 0.5 | 2 | 2 | | STAAU 13 | 0.06 | 2 | 2 |
| | STAAU 14 | 0.5 | 2 | 1 | | STAAU 14 | 0.5 | 16 | 16 |
| | STAAU 15 | 0.5 | 2 | 2 | | STAAU 15 | 0.25 | 2 | 1 |
| | STAAU 16 | 0.5 | 8 | 16 | | STAAU 16 | 0.25 | 2 | 2 |
| | STAAU 17 | 0.5 | 1 | 1 | | STAAU 17 | 0.12 | 1 | 2 |
| | STAAU 18 | 0.5 | 0.5 | 1 | | STAAU 18 | 0.12 | 2 | 2 |
| | STAAU 19 | 0.5 | 0.5 | 1 | | STAAU 19 | 0.12 | 2 | 2 |
| | STAAU 20 | 1 | 0.25 | 1 | | STAAU 20 | 0.12 | 2 | 2 |
| | STAAU 21 | 0.5 | 0.5 | 2 | | STAAU 21 | 0.5 | 4 | 2 |
| | STAAU 22 | 0.5 | 0.25 | 1 | | STAAU 22 | 0.25 | 2 | 4 |
| | STAAU 23 | 0.5 | 0.5 | 1 | | STAAU 23 | 0.25 | 2 | 4 |
| | STAAU 24 | 0.5 | 1 | 2 | | STAAU 24 | 0.25 | 16 | 8 |
| | STAAU 25 | 0.5 | 2 | 1 | | STAAU 25 | 0.12 | 2 | 1 |
| | STAAU 26 | 0.5 | 2 | 2 | | STAAU 26 | 0.12 | 2 | 2 |
| | STAAU 27 | 0.5 | 2 | 2 | | STAAU 27 | 0.06 | 2 | 4 |
| | STAAU 28 | 1 | 8 | 8 | | STAAU 28 | 0.25 | 2 | 4 |
| | STAAU 29 | 0.5 | 1 | 0.5 | | STAAU 29 | 0.06 | 2 | 2 |
| | STAAU 30 | 0.5 | 1 | 2 | | STAAU 30 | 0.25 | 2 | 4 |
| | | MIC Rep 1-3 | MBEC CC 1-3 | MBEC NBT 1-3 | | | MIC Rep 1-3 | MBEC CC 1-3 | MBEC NBT 1-3 |
| | QC 29213 | 1 | 2 | 2 | | QC 29213 | 0.25 | 2 | 2 |
| | QC 29213 | 1 | 1 | 1 | | QC 29213 | 0.25 | 2 | 4 |
| | QC 29213 | 1 | 2 | 2 | | QC 29213 | 0.25 | 2 | 4 |
| | | MIC Rep 4-6 | MBEC CC 4-6 | MBEC NBT 4-6 | | | MIC Rep 4-6 | MBEC CC 4-6 | MBEC NBT 4-6 |
| | QC 29213 | 1 | 2 | 2 | | QC 29213 | 0.5 | 2 | 2 |
| | QC 29213 | 1 | 2 | 2 | | QC 29213 | 0.25 | 2 | 2 |
| | QC 29213 | 1 | 2 | 2 | | QC 29213 | 0.25 | 2 | 2 |

Supplementary Table 3 – Mean MIC and MBEC E. coli vs Ciprofloxacin, Mean MIC and MBEC E.coli vs Piperacillin /Tazobactam

| | Organism No | mg/L | | |
|---|---|---|---|---|
| | | MIC | MBEC CC | MBEC NBT |
| Ciprofloxacin | ESCCO 1 | 0.008 | 0.12 | 0.06 |
| | ESCCO 2 | >1 | >1 | 1 |
| | ESCCO 3 | >1 | >1 | >1 |
| | ESCCO 4 | 0.008 | 0.12 | 0.03 |
| | ESCCO 5 | 0.004 | 0.008 | 0.008 |
| | ESCCO 6 | 0.25 | 0.25 | 0.25 |
| | ESCCO 7 | 0.008 | 0.008 | 0.008 |
| | ESCCO 8 | <0.002 | 0.004 | 0.008 |
| | ESCCO 9 | 0.25 | 0.5 | 0.25 |
| | ESCCO 10 | 0.015 | 0.12 | 0.25 |
| | ESCCO 11 | 0.015 | 0.03 | 0.25 |
| | ESCCO 12 | 0.008 | 0.015 | 0.015 |
| | ESCCO 13 | 0.008 | 0.004 | 0.004 |
| | ESCCO 14 | 0.25 | 0.25 | 0.25 |
| | ESCCO 15 | 0.25 | >1 | 1 |
| | ESCCO 16 | 0.25 | >1 | 1 |
| | ESCCO 17 | >1 | >1 | >1 |
| | ESCCO 18 | 0.015 | 0.06 | 0.03 |
| | ESCCO 19 | >1 | >1 | >1 |
| | ESCCO 20 | 0.015 | 0.015 | 0.015 |
| | ESCCO 21 | 0.12 | 0.5 | 0.5 |
| | ESCCO 22 | >1 | >1 | >1 |
| | ESCCO 23 | 0.008 | 0.25 | 1 |
| | ESCCO 24 | 0.008 | 0.03 | 0.015 |
| | ESCCO 25 | 0.008 | 0.008 | 0.008 |
| | ESCCO 26 | 0.008 | 0.008 | 0.015 |
| | ESCCO 27 | 0.004 | 0.004 | 0.008 |
| | ESCCO 28 | 0.015 | 0.015 | 0.03 |
| | ESCCO 29 | 0.015 | 0.015 | 0.015 |
| | ESCCO 30 | 0.015 | 0.008 | 0.015 |
| | | MIC Rep 1-3 | MBEC CC 1-3 | MBEC NBT 1-3 |
| | QC 25922 | 0.004 | 0.008 | 0.008 |
| | QC 25923 | 0.004 | 0.015 | 0.015 |
| | QC 25924 | 0.008 | 0.008 | 0.015 |
| | | MIC Rep 4-6 | MBEC CC 4-6 | MBEC NBT 4-6 |
| | QC 25922 | 0.008 | 0.015 | 0.15 |
| | QC 25923 | 0.008 | 0.008 | 0.008 |
| | QC 25924 | 0.008 | 0.008 | 0.008 |

| | Organism No | mg/L | | |
|---|---|---|---|---|
| | | MIC | MBEC CC | MBEC NBT |
| Piperacillin- Tazobactam | ESCCO 1 | 0.008 | 0.5 | 0.5 |
| | ESCCO 2 | 4 | 16 | 16 |
| | ESCCO 3 | 8 | 16 | >16 |
| | ESCCO 4 | 0.12 | 2 | 2 |
| | ESCCO 5 | 0.06 | 0.5 | 0.5 |
| | ESCCO 6 | 0.25 | 2 | 2 |
| | ESCCO 7 | 0.06 | 1 | 2 |
| | ESCCO 8 | 0.015 | 2 | 2 |
| | ESCCO 9 | 0.03 | 2 | 2 |
| | ESCCO 10 | 0.015 | 1 | 1 |
| | ESCCO 11 | 0.03 | 2 | 2 |
| | ESCCO 12 | 0.03 | 2 | 2 |
| | ESCCO 13 | 0.015 | 1 | 1 |
| | ESCCO 14 | 0.06 | 2 | 2 |
| | ESCCO 15 | 0.12 | 1 | 1 |
| | ESCCO 16 | 0.12 | 0.5 | 0.5 |
| | ESCCO 17 | 4 | >16 | >16 |
| | ESCCO 18 | 0.03 | 1 | 1 |
| | ESCCO 19 | 8 | 8 | 16 |
| | ESCCO 20 | 0.03 | 1 | 1 |
| | ESCCO 21 | 0.015 | 0.5 | 0.5 |
| | ESCCO 22 | 0.25 | 0.25 | 0.25 |
| | ESCCO 23 | 4 | 16 | 16 |
| | ESCCO 24 | 0.12 | 1 | 1 |
| | ESCCO 25 | 0.12 | 1 | 1 |
| | ESCCO 26 | 0.25 | 1 | 2 |
| | ESCCO 27 | 0.03 | 1 | 4 |
| | ESCCO 28 | 0.12 | 1 | 1 |
| | ESCCO 29 | 0.03 | 1 | 1 |
| | ESCCO 30 | 0.03 | 1 | 1 |
| | | MIC Rep 1-3 | MBEC CC 1-3 | MBEC NBT 1-3 |
| | QC 25922 | 4 | 16 | 16 |
| | QC 25923 | 4 | 8 | 8 |
| | QC 25924 | 4 | 16 | 8 |
| | | MIC Rep 4-6 | MBEC CC 4-6 | MBEC NBT 4-6 |
| | QC 25922 | 4 | 16 | 16 |
| | QC 25923 | 2 | 8 | 8 |
| | QC 25924 | 4 | 8 | 16 |

Supplementary Table 4 – Mean MIC and MBEC Pseudomonas aeruginosa vs ciprofloxacin, Mean MIC MBEC Pseudomonas aeruginosa vs Colistin

| | Organism | mg/L | | | | Organism | mg/L | | |
|---|---|---|---|---|---|---|---|---|---|
| | No | MIC | MBEC CC | MBEC NBT | | No | MIC | MBEC CC | MBEC NBT |
| Ciprofloxacin | PSEAE 1 | 0.12 | 0.25 | 0.5 | Colistin | PSEAE 1 | 0.5 | 2 | 1 |
| | PSEAE 2 | 0.12 | 0.25 | 0.5 | | PSEAE 2 | 0.06 | 0.5 | 2 |
| | PSEAE 3 | 0.06 | 0.12 | 0.25 | | PSEAE 3 | 0.12 | 1 | 0.5 |
| | PSEAE 4 | 0.06 | 0.25 | 0.5 | | PSEAE 4 | 0.12 | 0.5 | 0.5 |
| | PSEAE 5 | 0.12 | 0.5 | 0.5 | | PSEAE 5 | 0.25 | 0.5 | 0.5 |
| | PSEAE 6 | <0.008 | 0.12 | 0.12 | | PSEAE 6 | 0.06 | 1 | 2 |
| | PSEAE 7 | 0.12 | 0.06 | 0.06 | | PSEAE 7 | 0.25 | 0.5 | 0.5 |
| | PSEAE 8 | 0.12 | 0.12 | 0.5 | | PSEAE 8 | 0.25 | 4 | 2 |
| | PSEAE 9 | 0.12 | 0.06 | 0.25 | | PSEAE 9 | 0.12 | 0.25 | 0.5 |
| | PSEAE 10 | 0.06 | 0.06 | 0.06 | | PSEAE 10 | 0.25 | 0.25 | 1 |
| | PSEAE 11 | 0.25 | 0.25 | 0.25 | | PSEAE 11 | 0.5 | 0.5 | 0.5 |
| | PSEAE 12 | 0.06 | 0.12 | 0.12 | | PSEAE 12 | 0.5 | 8 | 4 |
| | PSEAE 13 | 0.12 | 0.06 | 0.12 | | PSEAE 13 | 0.25 | 0.25 | 0.5 |
| | PSEAE 14 | 0.12 | 0.06 | 0.25 | | PSEAE 14 | 0.12 | 0.25 | 0.25 |
| | PSEAE 15 | 0.06 | 0.12 | 0.25 | | PSEAE 15 | 0.5 | 1 | 2 |
| | PSEAE 16 | 0.12 | 0.12 | 0.5 | | PSEAE 16 | 0.25 | 0.5 | 0.5 |
| | PSEAE 17 | 0.5 | 2 | 4 | | PSEAE 17 | 2 | 4 | 4 |
| | PSEAE 18 | 0.12 | 0.25 | 0.12 | | PSEAE 18 | 0.12 | 0.12 | 0.5 |
| | PSEAE 19 | 0.06 | 0.5 | 0.5 | | PSEAE 19 | 0.25 | 0.5 | 1 |
| | PSEAE 20 | 0.12 | 0.5 | 0.5 | | PSEAE 20 | 0.25 | 0.5 | 0.5 |
| | PSEAE 21 | 0.06 | 0.5 | 0.25 | | PSEAE 21 | 0.12 | 0.25 | 0.5 |
| | PSEAE 22 | 0.06 | 0.12 | 1 | | PSEAE 22 | 0.12 | 2 | 2 |
| | PSEAE 23 | 8 | 8 | 16 | | PSEAE 23 | 4 | 16 | 16 |
| | PSEAE 24 | 32 | 128 | 128 | | PSEAE 24 | 8 | 256 | 128 |
| | PSEAE 25 | 128 | 128 | 256 | | PSEAE 25 | 8 | 256 | 256 |
| | PSEAE 26 | 0.12 | 0.25 | 0.5 | | PSEAE 26 | 0.12 | 1 | 2 |
| | PSEAE 27 | 8 | 256 | 256 | | PSEAE 27 | 4 | 128 | 128 |
| | PSEAE 28 | 64 | 64 | 64 | | PSEAE 28 | 16 | 64 | 128 |
| | PSEAE 29 | 0.25 | 0.5 | 0.5 | | PSEAE 29 | 0.12 | 0.25 | 0.25 |
| | PSEAE 30 | 0.06 | 1 | 0.5 | | PSEAE 30 | 0.06 | 0.5 | 0.5 |
| | | MIC Rep 1-3 | MBEC CC 1-3 | MBEC NBT 1-3 | | | MIC Rep 1-3 | MBEC CC 1-3 | MBEC NBT 1-3 |
| | QC 27853 | 0.25 | 2 | 2 | | QC 27853 | 1 | 4 | 4 |
| | QC 27853 | 0.25 | 1 | 1 | | QC 27853 | 1 | 2 | 4 |
| | QC 27853 | 0.12 | 2 | 4 | | QC 27853 | 1 | 4 | 4 |
| | | MIC Rep 4-6 | MBEC CC 4-6 | MBEC NBT 4-6 | | | MIC Rep 4-6 | MBEC CC 4-6 | MBEC NBT 4-6 |
| | QC 27853 | 0.12 | 2 | 2 | | QC 27853 | 1 | 4 | 4 |
| | QC 27853 | 0.12 | 2 | 4 | | QC 27853 | 0.5 | 4 | 4 |
| | QC 27853 | 0.12 | 2 | 2 | | QC 27853 | 1 | 4 | 8 |

Supplementary Table 5 - Mean MIC and MBEC Klebsiella pneumonia vs ciprofloxacin, Mean MIC and MBEC Klebsiella pneumonia vs Colistin

Ciprofloxacin

| Organism | mg/L | | |
|---|---|---|---|
| No | MIC | MBEC CC | MBEC NBT |
| KLEPN 1 | 0.25 | 0.5 | 0.5 |
| KLEPN 2 | 0.008 | 0.03 | 0.03 |
| KLEPN 3 | 0.008 | 0.5 | 0.25 |
| KLEPN 4 | 0.015 | 0.12 | 0.12 |
| KLEPN 5 | 0.015 | 0.25 | 0.25 |
| KLEPN 6 | 0.25 | 0.25 | 0.25 |
| KLEPN 7 | 1 | >32 | >32 |
| KLEPN 8 | 2 | 4 | 4 |
| KLEPN 9 | 2 | 8 | 4 |
| KLEPN 10 | 2 | 4 | 4 |
| KLEPN 11 | 1 | 16 | 16 |
| KLEPN 12 | 2 | 4 | 4 |
| KLEPN 13 | 0.5 | 4 | 4 |
| KLEPN 14 | 0.25 | 8 | 8 |
| KLEPN 15 | 4 | 4 | 4 |
| KLEPN 16 | 4 | 16 | 16 |
| KLEPN 17 | 0.25 | 2 | 4 |
| KLEPN 18 | 1 | 4 | 4 |
| KLEPN 19 | 1 | 8 | 8 |
| KLEPN 20 | 0.5 | 4 | 4 |
| KLEPN 21 | 8 | >32 | >32 |
| KLEPN 22 | 0.5 | 16 | 16 |
| KLEPN 23 | 4 | >32 | >32 |
| KLEPN 24 | 1 | 2 | 2 |
| KLEPN 25 | 4 | 32 | 16 |
| KLEPN 26 | 8 | 16 | 16 |
| KLEPN 27 | 4 | >32 | >32 |
| KLEPN 28 | 1 | 16 | 16 |
| KLEPN 29 | 0.03 | 0.5 | 1 |
| KLEPN 30 | 1 | 4 | 4 |
| KLEPN 31 | 4 | 8 | 8 |
| KLEPN 32 | 0.03 | 1 | 0.5 |
| | MIC Rep 1-3 | MBEC CC 1-3 | MBEC NBT 1-3 |
| QC 700603 | 0.5 | 4 | 4 |
| QC 700603 | 0.5 | 4 | 4 |
| QC 700603 | 0.5 | 4 | 4 |
| | MIC Rep 4-6 | MBEC CC 4-6 | MBEC NBT 4-6 |
| QC 700603 | 0.5 | 4 | 4 |
| QC 700603 | 0.5 | 4 | 4 |
| QC 700603 | 0.25 | 4 | 4 |

Colistin

| Organism | mg/L | | |
|---|---|---|---|
| No | MIC | MBEC CC | MBEC NBT |
| KLEPN 1 | 0.25 | 16 | 16 |
| KLEPN 2 | 0.25 | 0.5 | 0.5 |
| KLEPN 3 | 0.25 | 1 | 1 |
| KLEPN 4 | 0.25 | 4 | 4 |
| KLEPN 5 | 0.25 | 8 | 8 |
| KLEPN 6 | 0.25 | 0.5 | 0.5 |
| KLEPN 7 | 0.25 | 1 | 1 |
| KLEPN 8 | 0.25 | 4 | 8 |
| KLEPN 9 | 0.25 | 2 | 2 |
| KLEPN 10 | 0.25 | 16 | 16 |
| KLEPN 11 | 0.5 | 32 | 32 |
| KLEPN 12 | 0.25 | 8 | 8 |
| KLEPN 13 | 0.25 | 1 | 1 |
| KLEPN 14 | 0.5 | >128 | >128 |
| KLEPN 15 | 1 | 16 | 16 |
| KLEPN 16 | 0.5 | 4 | 4 |
| KLEPN 17 | 1 | 8 | 8 |
| KLEPN 18 | 0.5 | 0.5 | 0.5 |
| KLEPN 19 | 0.5 | 1 | 1 |
| KLEPN 20 | 0.25 | 4 | 4 |
| KLEPN 21 | 0.5 | >128 | >128 |
| KLEPN 22 | 0.5 | 64 | 64 |
| KLEPN 23 | 0.5 | 16 | 16 |
| KLEPN 24 | 0.25 | 1 | 1 |
| KLEPN 25 | 0.25 | 2 | 1 |
| KLEPN 26 | 0.25 | 4 | 4 |
| KLEPN 27 | 0.25 | 16 | 16 |
| KLEPN 28 | 0.25 | 8 | 8 |
| KLEPN 29 | 0.5 | >128 | >128 |
| KLEPN 30 | 0.5 | 8 | 8 |
| KLEPN 31 | 0.5 | 64 | 64 |
| KLEPN 32 | 0.25 | 2 | 2 |
| | MIC Rep 1-3 | MBEC CC 1-3 | MBEC NBT 1-3 |
| QC 700603 | 0.5 | 8 | 8 |
| QC 700603 | 1 | 4 | 8 |
| QC 700603 | 1 | 8 | 8 |
| | MIC Rep 4-6 | MBEC CC 4-6 | MBEC NBT 4-6 |
| QC 700603 | 1 | 8 | 8 |
| QC 700603 | 1 | 8 | 8 |
| QC 700603 | 1 | 8 | 8 |

Sup table 6 – CFU/mL raw data

| | | | | | | | | | | | | |
|---|---|---|---|---|---|---|---|---|---|---|---|---|
| | Planktonic | | | | | | | | | | | |
| | GC | | | X2 | | | X4 | | | X8 | | |
| 0 | 1800000 | 1300000 | 1900000 | 1700000 | 1200000 | 1100000 | 1000000 | 1300000 | 1800000 | 1400000 | 1500000 | 1600000 |
| 2 | 8700000 | 9200000 | 7700000 | 580000 | 720000 | 810000 | 620000 | 540000 | 710000 | 520000 | 470000 | 630000 |
| 4 | 1.8E+08 | 4E+08 | 4E+08 | 120000 | 220000 | 240000 | 95000 | 120000 | 550000 | 210000 | 180000 | 110000 |
| 8 | 4E+08 | 4E+08 | 4E+08 | 62000 | 74000 | 82000 | 36000 | 71000 | 28000 | 52000 | 93000 | 12000 |
| 24 | 4E+08 | 4E+08 | 4E+08 | 5600 | 1200 | 8200 | 72 | 9 | 9100 | 92 | 1500 | 1 |
| | Biofilm | | | | | | | | | | | |
| | GC | | | X2 | | | X4 | | | X8 | | |
| 0 | 1200000 | 1600000 | 1900000 | 1000000 | 1100000 | 890000 | 1300000 | 1700000 | 1200000 | 1600000 | 1900000 | 1900000 |
| 2 | 8200000 | 12000000 | 22000000 | 1500000 | 1800000 | 1100000 | 1100000 | 1500000 | 1000000 | 820000 | 910000 | 57000 |
| 4 | 55000000 | 62000000 | 82000000 | 3200000 | 4200000 | 1200000 | 720000 | 810000 | 73000 | 120000 | 136000 | 92000 |
| 8 | 1.2E+08 | 2.5E+08 | 4E+08 | 9200000 | 4800000 | 7100000 | 81000 | 110000 | 52000 | 82000 | 72000 | 51000 |
| 24 | 4E+08 | 4E+08 | 4E+08 | 12000000 | 5200000 | 9100000 | 53000 | 320000 | 130000 | 650 | 12000 | 180000 |
| | KLEPN | vs | CIPRO | | | | | | | | | |
| | Planktonic | | | | | | | | | | | |
| | GC | | | X2 | | | X4 | | | X8 | | |
| 0 | 1600000 | 1800000 | 2100000 | 1300000 | 1700000 | 1200000 | 1900000 | 1700000 | 1000000 | 1500000 | 1500000 | 1900000 |
| 2 | 8200000 | 9200000 | 4700000 | 820000 | 910000 | 900000 | 1100000 | 720000 | 620000 | 120000 | 380000 | 170000 |
| 4 | 4E+08 | 4E+08 | 32000000 | 53000 | 74000 | 81000 | 28000 | 36000 | 32000 | 16000 | 22000 | 72000 |
| 8 | 4E+08 | 4E+08 | 4E+08 | 61000 | 52000 | 74000 | 310000 | 110000 | 130000 | 8200 | 6300 | 7100 |
| 24 | 4E+08 | 3.8E+08 | 4E+08 | 5200000 | 4200000 | 3800000 | 590000 | 1400 | 360000 | 25 | 5200 | 14000 |
| | Biofilm | | | | | | | | | | | |
| | GC | | | X2 | | | X4 | | | X8 | | |
| 0 | 2300000 | 1700000 | 1500000 | 1200000 | 1900000 | 1800000 | 1700000 | 1500000 | 1200000 | 960000 | 1400000 | 1500000 |
| 2 | 5900000 | 12000000 | 7200000 | 2500000 | 1800000 | 2000000 | 2200000 | 2900000 | 1300000 | 1600000 | 1200000 | 1300000 |
| 4 | 20000000 | 32000000 | 17000000 | 5200000 | 6700000 | 4200000 | 3200000 | 4100000 | 2000000 | 1000000 | 960000 | 1100000 |
| 8 | 4E+08 | 4E+08 | 4E+08 | 12000000 | 22000000 | 9200000 | 3100000 | 5200000 | 2200000 | 880000 | 910000 | 860000 |
| 24 | 4E+08 | 4E+08 | 4E+08 | 15000000 | 30000000 | 14000000 | 3200000 | 7200000 | 2800000 | 120000 | 7200 | 820000 |
| | STRPN | vs | LEVO | | | | | | | | | |
| | Planktonic | | | | | | | | | | | |
| | GC | | | X2 | | | X4 | | | X8 | | |
| 0 | 1600000 | 1700000 | 1200000 | 1800000 | 1600000 | 1200000 | 1200000 | 1900000 | 2100000 | 1500000 | 1000000 | 1400000 |
| 2 | 6200000 | 7100000 | 5600000 | 1100000 | 880000 | 720000 | 860000 | 780000 | 1600000 | 620000 | 710000 | 530000 |
| 4 | 9100000 | 9800000 | 11000000 | 520000 | 690000 | 620000 | 320000 | 120000 | 850000 | 120000 | 63000 | 77000 |
| 8 | 25000000 | 32000000 | 4E+08 | 32000 | 52000 | 61000 | 150000 | 22000 | 27000 | 9900 | 14000 | 920 |
| 24 | 4E+08 | 4E+08 | 4E+08 | 18000000 | 2700000 | 2100000 | 610000 | 120000 | 240000 | 3200 | 2800 | 17 |
| | Biofilm | | | | | | | | | | | |
| | GC | | | X2 | | | X4 | | | X8 | | |
| 0 | 1800000 | 950000 | 890000 | 1100000 | 960000 | 1400000 | 870000 | 1300000 | 1600000 | 1100000 | 1000000 | 890000 |
| 2 | 12000000 | 8200000 | 9200000 | 1500000 | 3200000 | 2800000 | 770000 | 670000 | 660000 | 1600000 | 890000 | 710000 |
| 4 | 95000000 | 4E+08 | 4E+08 | 82000000 | 66000000 | 91000000 | 12000000 | 22000000 | 9200000 | 620000 | 710000 | 630000 |
| 8 | 4E+08 | 4E+08 | 4E+08 | 4E+08 | 4E+08 | 4E+08 | 20000000 | 36000000 | 10000000 | 730000 | 410000 | 260000 |
| 24 | 4E+08 | 4E+08 | 4E+08 | 4E+08 | 4E+08 | 4E+08 | 17000000 | 62000000 | 11000000 | 52000 | 62000 | 87000 |

Supplementary figure 1 – Typical challenge plate

| | 0.12 | 0.25 | 0.5 | 1 | 2 | 4 | 8 | 16 | 32 | 64 | GC | SC | MIC mg/L |
|---|---|---|---|---|---|---|---|---|---|---|---|---|---|
| A | | | | | | | | | | | | | |
| B | | | | | | | | | | | | | |
| C | | | | | | | | | | | | | |
| D | | | | | | | | | | | | | |
| E | | | | | | | | | | | | | |
| F | | | | | | | | | | | | | |
| G | | | | | | | | | | | | | |
| H | | | | | | | | | | | | | |
